# Genome-wide CRISPRi maps pneumococcal attachment and intracellular survival during single infection and Influenza co-infection

**DOI:** 10.64898/2026.09.18.752567

**Authors:** Philipp DK. Walch, Petr Broz, Jan-Willem Veening

## Abstract

*Streptococcus pneumoniae* remains a leading cause of bacterial pneumonia and frequently causes severe secondary infections following influenza A virus (IAV) infection. Yet, the genetic requirements underlying pneumococcal pathogenicity during viral co-infection remain incompletely understood. Here, we employed an inducible genome-wide CRISPR interference (CRISPRi) library to systematically investigate pneumococcal gene fitness during epithelial attachment and intracellular survival in macrophages under both single-infection and IAV co-infection conditions.

Screening across multiple host cell types and viral strains identified 42 pneumococcal genes affecting host-cell attachment and 63 genes influencing intracellular survival. While global patterns of gene fitness were largely conserved between single infection and co-infection, genes involved in cell envelope biogenesis, including the undecaprenyl pyrophosphate phosphatase *uppP*, became increasingly important during co-infection. Pharmacological inhibition of cell wall biosynthesis recapitulated several genetic phenotypes and revealed enhanced sensitivity to bacitracin in virally altered host environments. Integration of CRISPRi screening with untargeted metabolomics uncovered extensive remodeling of nucleotide metabolism during infection. Disruption of nucleoside transport altered bacterial fitness, morphology, capsule expression, and host-cell attachment. Uridine availability emerged as a key regulator linking metabolic adaptation to virulence-associated traits, highlighting a trade-off between bacterial growth and adherence.

Taken together, our study provides a genome-scale resource of pneumococcal fitness determinants during infection and demonstrates that influenza co-infection selectively reshapes bacterial genetic dependencies rather than globally altering pathogenicity programs. These findings identify cell envelope homeostasis and nucleotide metabolism as central regulators of pneumococcal virulence and reveal potential targets for future antimicrobial and anti-virulence strategies.

## Introduction

*Streptococcus pneumoniae* remains one of the leading causes of bacterial pneumonia^1,2^, meningitis^3,4^ and sepsis^2,5^ worldwide, resulting in substantial morbidity and mortality across both pediatric and adult populations^6^. Despite the implementation of effective conjugate vaccines, pneumococcal disease continues to pose a major global health burden, partly due to serotype replacement and the emergence of strains with reduced susceptibility to multiple antimicrobial classes^7,8^. Antimicrobial resistance (AMR) in *S. pneumoniae* increasingly compromises treatment efficacy and giving it priority status in global AMR surveillance and research initiatives, including the World Health Organization^9,10^. This underscores the urgent need to identify new molecular targets that can be exploited for antimicrobial or anti-infective intervention.

The challenge posed by pneumococcal infections is further amplified during respiratory viral co-infections^11^. Among these, influenza A virus (IAV) represents the most clinically relevant viral partner of *S. pneumoniae*^12–14^. Historical analyses of seasonal and pandemic influenza outbreaks have consistently demonstrated that secondary bacterial infections contribute substantially to disease severity, hospitalization, and mortality^15^. Clinical studies have reported frequent pneumococcal co-detection in patients with severe influenza^16,17^, and meta-analyses of hospitalized influenza cohorts identify *S. pneumoniae* as one of the most prevalent bacterial co-occurring pathogens^18,19^. These observations reflect a well-established biological synergy in which influenza-induced epithelial damage, altered innate immunity, depletion of alveolar macrophages and changes in the respiratory microenvironment facilitate pneumococcal colonization and invasion^20–22^. Additionally, there is evidence of direct interkingdom interaction that affects pathogenicity during co-occurrence^23^.

Recent advances in functional bacterial genomics have made it possible to systematically investigate gene fitness across diverse environmental and host-associated conditions^24–26^. In particular, CRISPR-interference (CRISPRi) has emerged as a powerful tool for genome-scale phenotypic screening by enabling inducible and tunable repression of target genes without permanently altering the genome^25,27,28^. Unlike transposon-based approaches, which cannot readily be employed to interrogate essential genes, CRISPRi permits graded knockdown of both essential and non-essential loci^29^. This capability enables the identification of essential genes whose importance varies according to environmental context^30^, genetic background^31^, or host niche^32^. Consequently, CRISPRi-based screening provides a comprehensive framework for dissecting bacterial fitness determinants and uncovering therapeutic targets that have so far remained inaccessible.

In this study, we employed an inducible genome-wide CRISPRi screening strategy to identify pneumococcal genes that are required for pathogenicity during both single infection and co-infection with distinct influenza A virus strains. Using bacterial attachment to host epithelial cells and intracellular survival within macrophages as quantitative readouts, we sought to identify context-dependent effects of gene repression on pneumococcal fitness during single infection and influenza co-infection. Furthermore, we investigated how influenza co-infection reshapes fitness patterns, revealing molecular interactions between host pathways and the two co-occurring pathogens. By identifying genetic dependencies that emerge specifically during co-infection, our work provides new insights into polymicrobial disease biology and highlights potential intervention points that may be leveraged to disrupt host-pathogen-pathogen interactions for the development of novel anti-infective strategies.

## Results

### Superinfection workflow and analysis pipeline produces high-quality data

During infection, pathogens interact with several host cells: Epithelial cells represent the first barrier for infecting pathogens, and successful attachment is essential to colonize and establish infection^33,34^. Subsequently, the host’s innate immune response is initiated. Macrophages represent a key cell type that is deployed to control pathogen spread^35^, and surviving clearance by macrophages is a second important aspect of pathogenicity^36^. We therefore used epithelial attachment and intracellular survival in macrophages as complementary readouts of pneumococcal pathogenicity.

To identify bacterial genes required for colonization and intracellular survival, we established a CRISPRi-seq infection workflow using two established *in vitro* model systems: A549 epithelial cells (attachment) and PMA-differentiated THP-1 macrophages (intracellular survival). Host cells were either left uninfected or pre-infected with different IAV strains: *Wyoming* (WY), a H3N2 strain^37^ and *Netherlands* (N), a H1N1 strain^38^, at a multiplicity of infection (MOI) of 2 for 4h or 24h. Cells were subsequently infected with a pre-induced genome-wide CRISPRi-library of an unencapsulated *S. pneumoniae* serotype 2 D39V strain, containing 1498 unique sgRNAs targeting all known operons in *S. pneumoniae* D39V^25,31^. Each clone in the library expresses a unique sgRNA that is constitutively expressed while the *dcas9* in under control of the IPTG-inducible P*lac* promoter^39^. Attached bacteria were recovered after 3h (Figure 1A i) and sgRNA abundance was quantified by sequencing. In THP-1 infection, extracellular bacteria were eliminated using recombinant LytA autolysin, which digests the pneumococcal cell wall^40^, before recovery of intracellular populations at later timepoints (Figure 1A ii).

**Figure 1:**
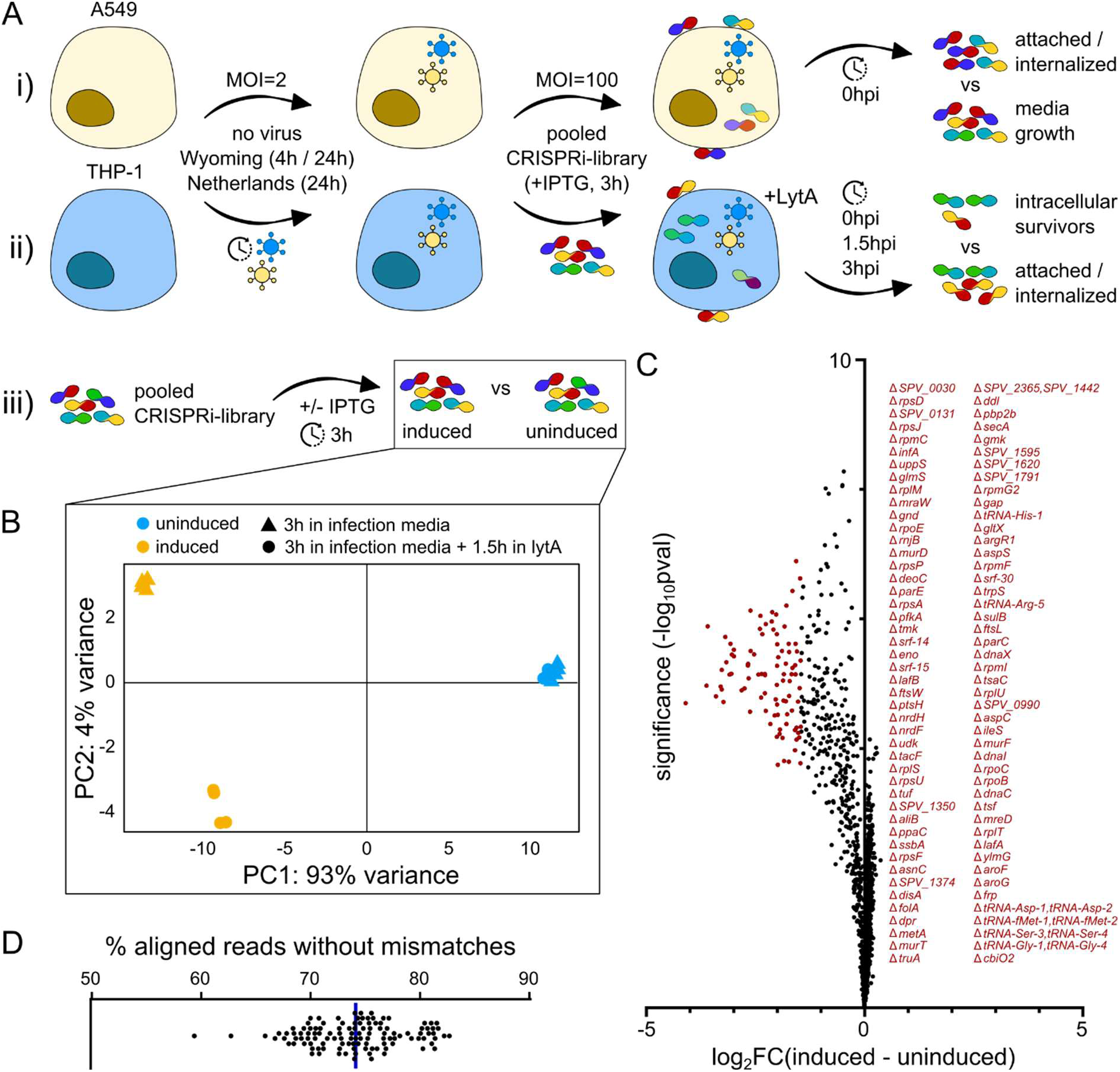
Workflow and general quality control. A) Schematic depicting the workflow of the CRISPRi screening: A549 (i) and THP-1 cells (ii) were infected with different IAV strains, as indicated, including an uninfected control. Pre-infected cells were then exposed to the previously induced *S. pneumoniae* library in the presence of IPTG for 3h. Subsequently, bacteria were harvested by washing and host cell lysis at the indicated timepoints. Additionally, an induction control (iii) was performed separately. B) PCA plot showing the discernability of induced and uninduced samples (bacterial growth in infection media). C) Volcano plot depicting the logarithmic fold change of barcode frequency in induced vs uninduced samples. Operons that are enriched on the left (indicated in red) are present at a lower abundance after induction, indicating that CRISPRi depletion has a fitness effect. D) Beeswarm plot that shows the distribution of aligned reads (without mismatches) after sequencing. Each dot represents an individual replicate per condition and timepoint.

We next assessed data quality and screen performance. Principal component analysis (PCA) revealed a clear separation between IPTG-induced (i.e. with *dcas9* expression) and uninduced bacteria (Figure 1A iii), confirming effective CRISPRi (Figure 1B). Consistent with this, depletion analysis recapitulated known fitness-associated genes that were underrepresented upon IPTG induction (Figure 1C, genes listed in red). We further demonstrated robust library representation across all infection conditions, with a high proportion of aligned reads and broad coverage across the genome (Figure 1D, Figure S1A). Importantly, neither viral co-infection nor later sampling timepoints introduced substantial global biases, and barcode counts remained sufficient across all experimental conditions (Figure S1B-D). Together, these analyses demonstrate that the workflow produces reproducible, high-quality datasets suitable for identifying infection-specific fitness determinants.

### A genome-wide screen identifies 42 genes that affect *S. pneumoniae* attachment

The established CRISPRi-seq infection workflow allowed us to systematically assess the contribution of individual pneumococcal genes to host-cell attachment. By comparing sgRNA abundance after 3h of attachment to the baseline library, we identified 42 genes whose depletion significantly altered attachment in at least one infection condition (21 genes increasing and 21 genes decreasing attachment, Figure 2A). This represents 2.80% of the genes assayed in the screen and is consistent with the limited global variance observed by PCA (Figure S2A). Attachment profiles are comparable between A549 and THP-1 cells, and most strongly enriched and depleted genes, including *SPV_2019-21*, the regulators of the adhesive factor CbpA^41^, are conserved (Figure 2B). This indicates a pathogen-driven attachment and invasion mechanism, rather than a dominating impact by the host.

**Figure 2:**
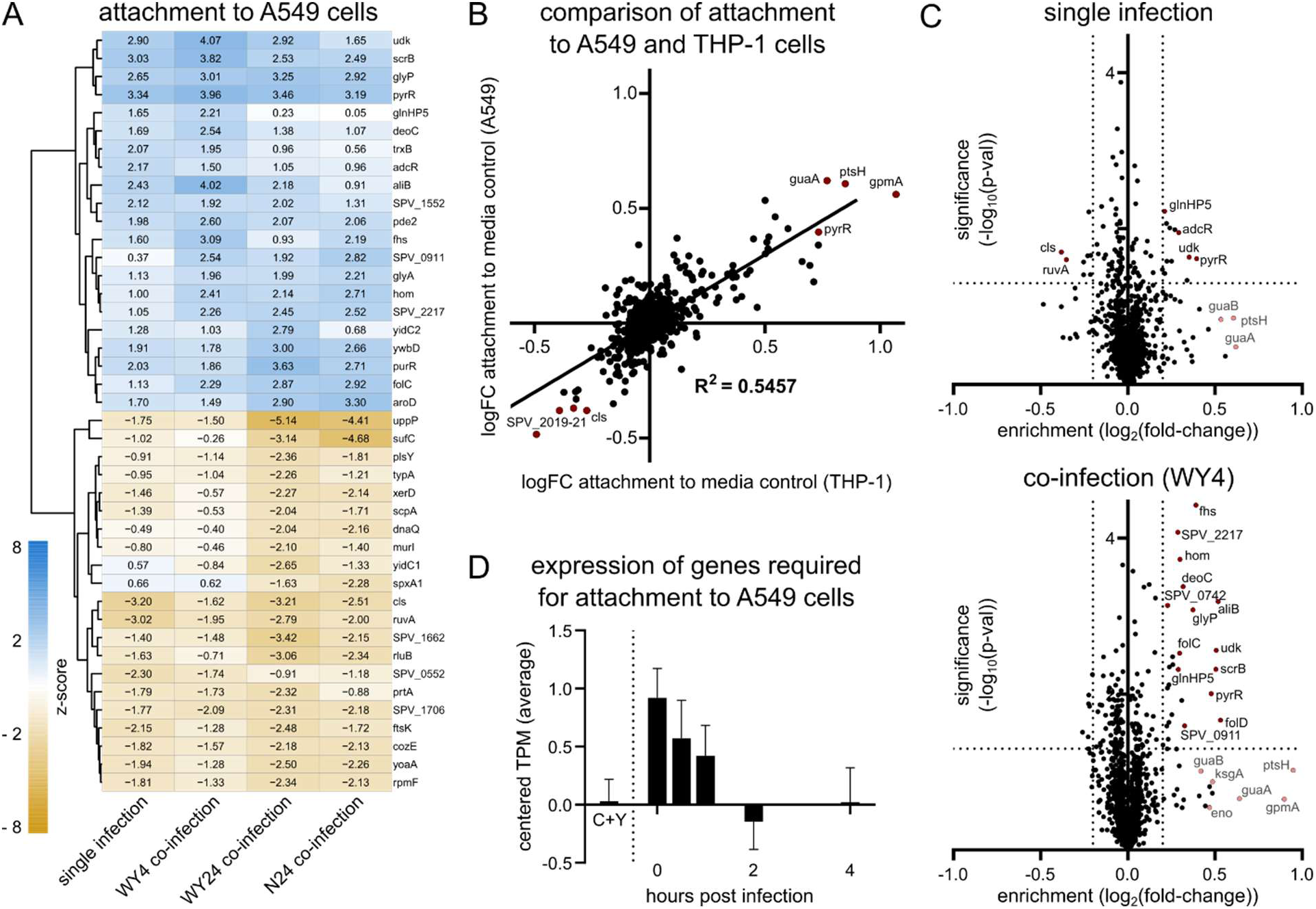
*S. pneumoniae* genes altering attachment to host cells. A) Reduced heatmap (z-score > 2 in at least one infection condition) of candidate genes which alter attachment when depleted. Shading according to z-score, as indicated in the legend (shades of blue: improved fitness upon deletion, shades of orange: gene required for fitness). Calculations were done with respect to the media control (fold change) and subsequently z-transformed. B) Scatter plots of log-fold changes of attachment to host cells (single infection) with respect to the media control. Each dot represents one gene, and candidates with highest and lowest enrichments are indicated by name. Pearson correlation between A549 and THP-1 is indicated. C) Two volcano plots showing logarithmic fold changes of barcodes detected after single infection (top panel) and *Wyoming* strain 4h co-infection (bottom panel) in attached bacteria compared to media growth. A subset of enriched or depleted genes are indicated by name. D) Genes that were essential for attachment (z-score < −2 in at least one condition) were analyzed with respect to their expression during infection ^42^. Bar graph depicts averages and standard deviation of centered TPM at each of the timepoints for which data was available (0h, 30min, 1h, 2, and 4h), as well as C+Y media as control.

The overall pattern of attachment-associated fitness was similar between single infection and IAV co-infection (Figure S2B, Figure 2A). However, viral pre-infection altered the attachment phenotypes of individual CRISPRi knockdown strains. Strains with reduced expression of glycolytic genes, including *eno*, *gmpA*, *gap* and *pyk*, showed a trend toward greater attachment during co-infection than during single infection, although these differences were not statistically significant (Figure 2C). Increased attachment during co-infection was also observed for knockdowns targeting genes involved in purine and pyrimidine metabolism and regulation, including *pyrR*. Pre-infection with the Wyoming IAV strain accentuated several knockdown phenotypes: it further reduced attachment-associated fitness in *uppP* and *sufC* knockdowns (Figure 2A) and further increased it in knockdowns that already showed enhanced attachment-associated fitness, as indicated by positive Z-scores (Figure 2C, Figure S2C).

To further assess the biological relevance of the identified candidates, we compared attachment-essential genes with publicly available infection transcriptomes^42^. Genes classified as important for attachment were predominantly upregulated during the early stages of infection (Figure 2D), supporting a role during host interaction. Additionally, we performed co-expression analysis (Figure S2D) to determine if specific clusters of co-expressed genes were systematically altering attachment behavior, yet no such cluster could be defined (Figure S2E). This indicates that expression changes are subtle and co-expression in non-infection conditions do not serve as reliable predictors for pathogenicity.

### CRISPRi-seq identifies 63 genes involved in *S. pneumoniae* survival within host macrophages

While *S. pneumoniae* is generally considered an extracellular pathogen, there are reports that describe survival inside macrophages of the host during infection^43,44^. We therefore investigated the impact of bacterial CRISPRi on the capability of *S. pneumoniae* to survive inside this reservoir. We identified 19 operons that reduced intracellular survival upon CRISPRi knockdown and 44 operons with improved survival (Figure 3A). This represents 4.21% of all genes assayed in the screening. We further assessed fold-changes in each individual condition (Figure S3A), as well as variance of sgRNA fingerprints (Figure S3B). In line with this, we also saw correlation between single and co-infection (Figure S3C), with shorter times of pre-infection yielding a higher correlation, indicating that viral pre-infection alters gene essentiality in a subtle and time-dependent manner.

**Figure 3:**
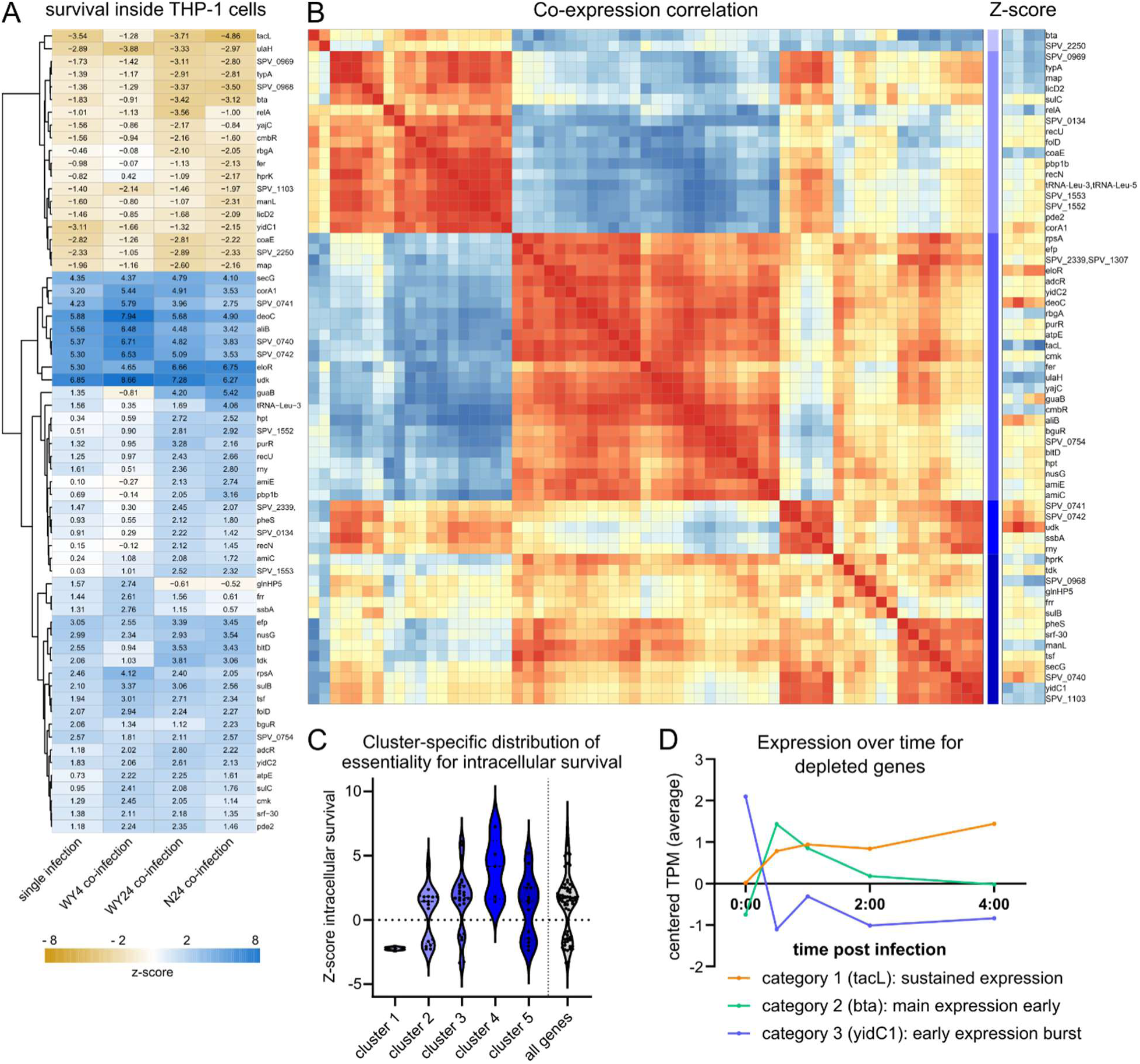
*S. pneumoniae* genes altering intracellular survival in macrophages. A) Reduced heatmap (z-score >2 in at least one infection condition) of candidate genes which alter intracellular survival when depleted. Shading according to z-score, as indicated in the legend. Calculations were done with respect to the 0hpi timepoint (fold change) and subsequently z-transformed. B) Matrix of co-expression correlation for all genes that significantly alter intracellular survival when targeted with CRISPRi. For each candidate gene, co-expression with all *S. pneumoniae* genes^42^ was assessed and converted into a co-expression fingerprint. Subsequently, pairwise correlations of fingerprints were calculated, and are depicted in the heatmap. Perfect correlation (score = 1) is colored in red, perfect anti-correlation (score = −1) in blue. K-means clustering (k=5) was performed to group candidate genes into clusters with similar fingerprints. C) Violin-plots for each of the clusters calculated in B: each dot represents one candidate gene, and displays the z-score in the screening. D) Expression profile during infection^42^ for three different groups of candidate genes that are essential for intracellular survival.

We selected genes whose repression significantly altered pneumococcal survival within macrophages in the CRISPRi screen. Using published transcriptomic data^42^, we then compared their genome-wide co-expression profiles and applied unsupervised k-means clustering to identify groups of genes with similar co-expression patterns and potentially related functions (Figure 3B). Of the five identified clusters, three (clusters 2, 3 and 5) did not show enrichment of genes specifically increasing or decreasing intracellular survival. Conversely, cluster 1 consisted of genes that were predominantly important for survival: *bta* (a bacteriocin transport accessory protein) and *SPV_2250* (a putative transcriptional regulator). Cluster 4, which is comprised of uridine-related genes and transporters, was enriched for genes that reduce fitness upon depletion (Figure 3C).

Focusing on all genes that are required for intracellular survival, we identified an increased expression during infection. We categorized genes into three groups using publicly available expression data during infection: a sustained expression during infection (Figure 3D, category 1), which included, among others, *tacL*, *rbgA* or *hprK*; an expression that occurred in early timepoints after infection (category 2), such as *bta*, *ulaH* or *manL*; and a boost immediately at the beginning of the infection process (category 3), such as *yidC1*, *map* or *SPV_0968* and *SPV_0969*.

### Experimental validation confirms essential, redundant, and neutral gene phenotypes under both infection conditions

To validate the CRISPRi-seq results, we created clean replacement-deletion mutants and assessed attachment or intracellular survival using colony forming unit (CFU) quantification. Candidate genes were selected to be representative across cell types, effect-size and direction observed in the screen. Deletions were introduced into the encapsulated D39V wildtype background rather than the Δ*cps* screening strain to provide an orthogonal validation system (Figure S4A). To account for potential growth-related effects, attachment was normalized to growth in media (Figure S4B), intracellular survival phenotypes were normalized to bacterial abundance at 0 hpi (Figure S4C).

Overall, the validation experiments supported the robustness of the screening dataset, with 76.9% of tested phenotypes reproducing across conditions (Figure 4A). Strong attachment and intracellular-survival phenotypes identified by CRISPRi (e.g. Δ*cls*, Δ*guaA*, Δ*tacL*) were readily confirmed in the corresponding deletion mutants (Figure 4B-C). For candidates with lower z-scores, the directionality of the phenotype was maintained but did not reach statistical significance (e.g. Δ*deoC*). Conversely, a small number of candidates could not be validated, particularly genes that altered bacterial entry into macrophages, which can affect the quantification of intracellular survival.

**Figure 4:**
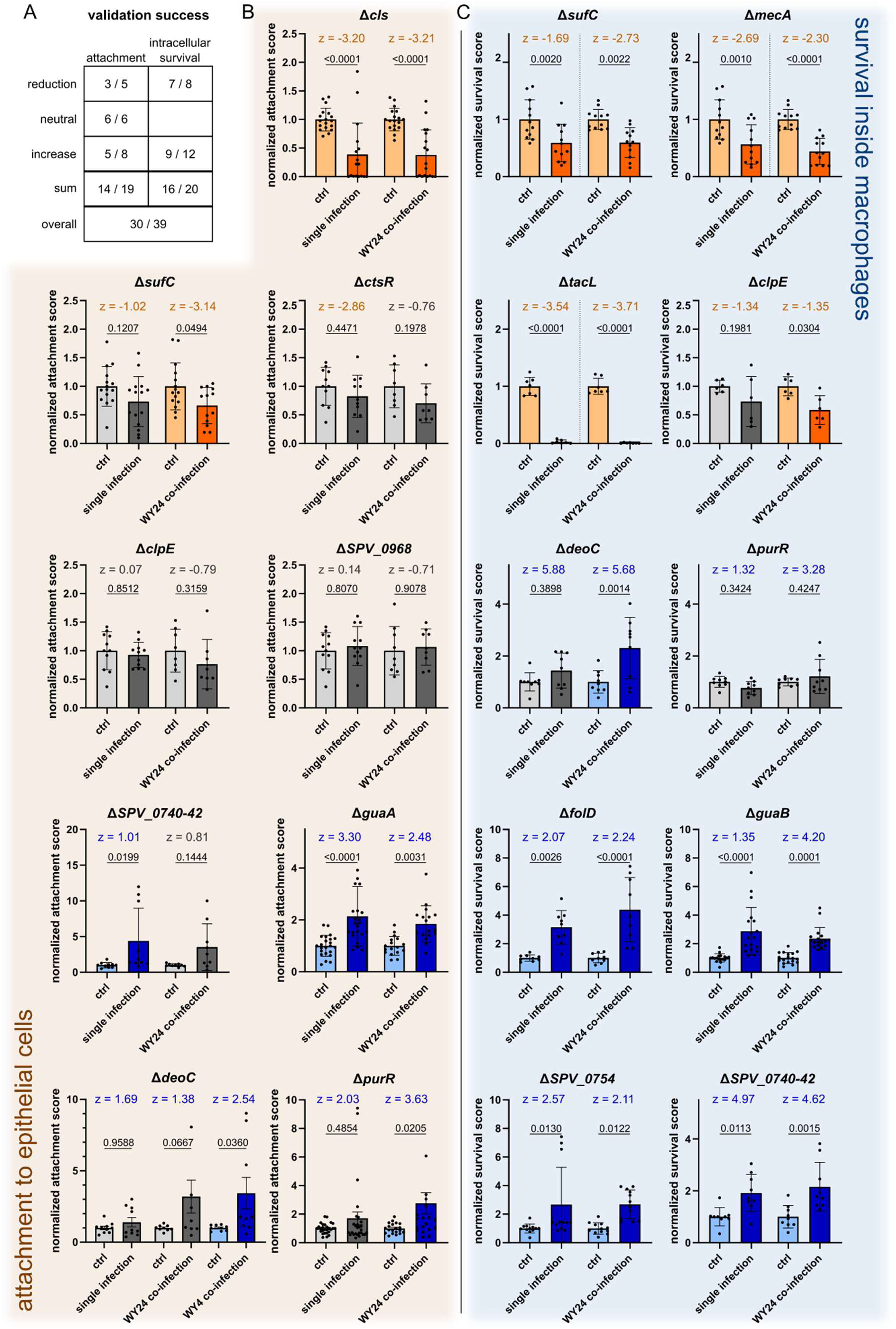
Validation using clean KO-mutant strains. A) Overview of validation success by assessed condition and directionality. B) Bar charts depicting the validation of attachment effects: Each dot represents one replicate. Attachment scores were calculated with respect to the media control, which is normalized to 1. Bars represent mean with standard deviation as error bars. One-Way-ANOVA was performed to calculate p-values. Additionally, z-scores are indicated for each assessed condition. Color coding as follows: Orange: Genes that are essential for attachment, Grey: Neutral genes, Blue: Genes that improve attachment when depleted by CRISPRi. C) As in B, but for intracellular survival. Normalization was performed with respect to the 0hpi timepoint.

In summary, these findings demonstrate that the major phenotypes identified in the CRISPRi screen can be reproduced in an independent genetic background and experimental framework. The validated candidates highlight previously implicated processes in pneumococcal pathogenicity, including cell-envelope homeostasis (*cls*, *tacL*) and metabolic regulation (*guaA*, *folD*, *purR*). It is noteworthy that targeting of single genes by CRISPRi, or their deletion, is limited in effect size by genetic redundancy^45–47^. This warrants the investigation of gene essentiality through different assays that span functional genetics, biochemical inhibition and combinatorial approaches^47,48^.

### Comparison to *in vivo* data highlights the importance of *uppP* in determining fitness during co-infection

To obtain a global overview of the identified fitness determinants, we first compared the attachment and intracellular-survival datasets and performed GO-term enrichment analysis. Several genes contributed to both aspects of pathogenicity, including *tacL*, *sufC* and *uppP*, indicating broader roles at the host-pathogen interface (Figure S5A). Likewise, genes whose depletion enhanced attachment frequently also improved intracellular survival, including *udk*, *pyrR* and *deoC*. GO-term enrichment revealed three major functional categories: biosynthesis of organic compounds, metabolic processes and protein transport (Figure 5A). Most enriched terms were associated with both attachment and intracellular survival (Figure S5B), suggesting shared biological requirements across infection stages.

**Figure 5:**
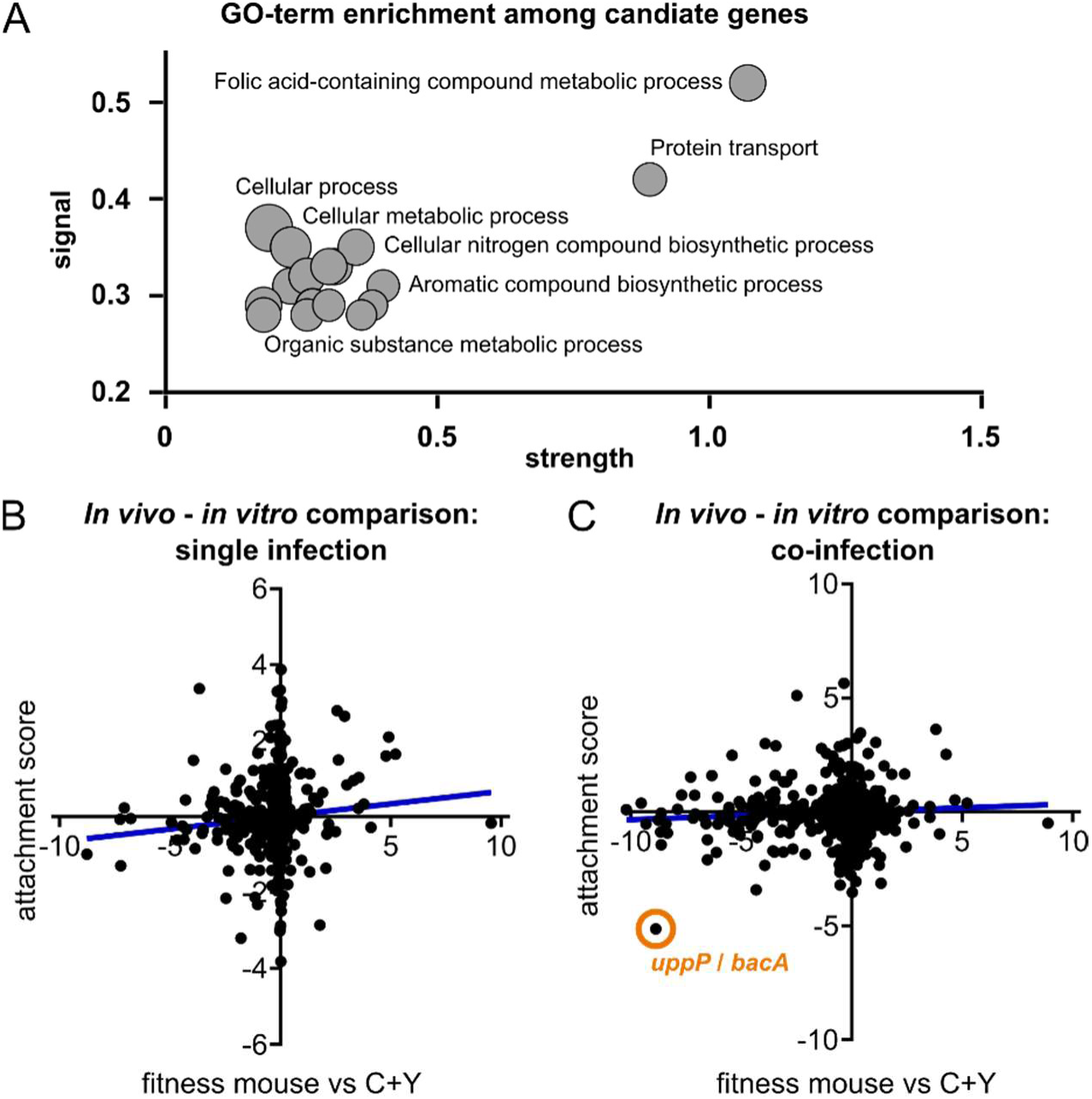
Global data analysis and comparison to existing datasets. A) GO-term enrichment of all genes that were significantly altered in either attachment of intracellular survival by CRISPRi. Analysis was done using String version 12.0^51–53^. Strength refers to the enrichment (log_10_(observed/expected), signal describes the weighted harmonic mean between the observed/expected-ratio and negative logarithmic false discovery rate (FDR). B) Correlation of attachment scores in this screen (y-axis) to score determined by CRISPRi during infection *in vivo*^49^ (single infection). Linear regression is indicated by the blue line C) As in B, but for IAV-co-infection. Strongest overlapping hit (*uppP*) is indicated by the orange circle.

We next compared our *in vitro* datasets with a recently published, genome-wide CRISPRi screen performed during pneumococcal infection and IAV co-infection *in vivo*^49^. This study used an encapsulated D39V background and therefore identified capsule-associated genes as dominant contributors to fitness. We sought to identify additional pathways emerging in a capsule-independent manner. During single infection, attachment scores displayed a weak but significant correlation with *in vivo* fitness values (Figure 5B), whereas intracellular-survival phenotypes showed no overall correlation (Figure S5C). Similar results were observed during co-infection, where no global correlation was detected (Figure 5C, Figure S5C).

Despite this limited overlap, which could be attributed to the presence of capsule on the infecting bacteria, one candidate emerged consistently across datasets. *uppP* was required for pathogenicity *in vivo*^49,50^ and strongly contributed to attachment during co-infection *in vitro* (Figure 5C). UppP catalyzes the dephosphorylation of undecaprenyl diphosphate, and is a crucial enzyme in the cell wall biogenesis of *S. pneumoniae*^50^. Together, these analyses highlight the context-dependent nature of pneumococcal fitness while identifying cell-envelope homeostasis, and particularly *uppP*, as a robust determinant of pathogenicity across infection models.

### Loss of *uppP* impairs *S. pneumoniae* fitness during infection, particularly during IAV co-infection

The previous experiments and global analyses suggest that the bacterial cell wall is a crucial determinant of host cell attachment. Therefore, we focused on cell wall associated genes with a fitness effect (Figure 6A). Several of those are antibiotic targets: *murA-1*, *murA-2*, which are inhibited by fosfomycin^54^, penicillin-binding proteins (PBPs), which are targets of amoxicillin and penicillin^55^, and *uppP* (*bacA*), which is mediating bacitracin-resistance^50,56,57^.

**Figure 6:**
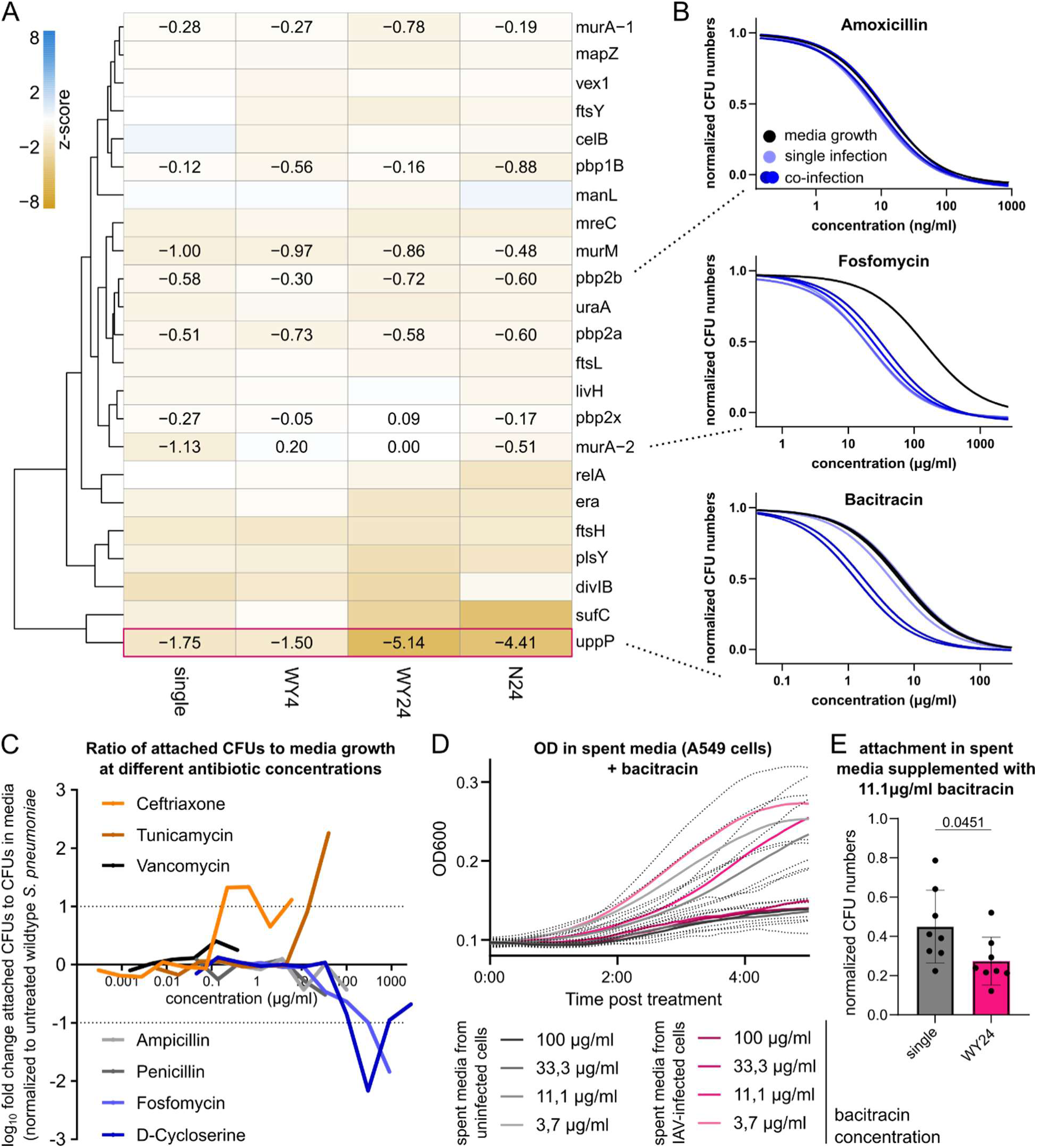
Chemical targeting of cell wall biosynthesis affects bacterial attachment. A) Reduced heatmap of the attachment dataset, focusing on cell wall related genes (by GO-term affiliation). Coloring and layout as in Figure 2. For known targets of antibiotics that were used in the study, z-scores are indicated. Pink frame highlights *uppP*, which displayed an increased fitness effect during co-infection (see Figure 2). B) Dose-response curves for three antibiotics with different mechanisms of action and different interaction during single and co-infection. Coloring as indicated in the legend. C) Ratio of attached CFUs to CFUs recovered from media during treatment with the indicated antibiotics at different concentrations. Values are normalized to the ratio in the untreated control. D) OD_600_ growth curves during treatment with bacitracin in spent media of uninfected or IAV-infected host cells. Shading of the solid line corresponds to the bacitracin concentration as indicated. Dotted line represents standard deviation intervals. E) CFUs (normalized to untreated control) recovered from cells after attachment in the presence of 11.1μg/ml bacitracin in spent media of uninfected or IAV-infected host cells. Each dot represents one replicate, and p-value was calculated by Students T-test with Welsh correction.

To test the targetability of these candidates, we treated *S. pneumoniae* with sublethal concentrations of a panel of antibiotics, comparing the dose-dependent inhibition of growth to that of attachment (Figure 6B): Amoxicillin and penicillin impacted attachment through growth inhibition (Figure S6A), fosfomycin treatment reduced attachment (IC_50_ = 35.7μg/ml for single infection) at much lower concentrations than growth (IC_50_ = 147.7μg/ml). Bacitracin did not impact attachment in single infection (IC_50_ = 6.1μg/ml, IC_50_ in media = 6.7μg/ml) but did reduce the required inhibitory concentration specifically during co-infection (IC_50_ = 0.21μg/ml).

Fosfomycin and cycloserine-D act in the cytoplasm upstream of the formation of Lipid-I^58,59^, and both reduced the fraction of attached bacteria compared to growth in media (Figure 6C). Other cell wall targeting drugs, specifically those targeting PBPs, either had a neutral effect or, as for ceftriaxone, increased the fraction of CFUs recovered after attachment. Intriguingly, a similar effect could be seen for tunicamycin, which induces ER stress in the host^60^ and perturbs Lipid-I and teichoic acid biosynthesis in bacteria^61,62^ (Figure 6C). By assessing cytotoxic effects during antibiotic treatment, we were able to exclude excessive host cell death as confounding factor (Figure S6B).

Finally, we investigated the molecular basis for the synergistic interaction between viral infection and bacitracin treatment. Since *uppP* is much more required for pneumococcal survival during co-infection (Figure 6A), mirroring the effect of bacitracin (Figure 6B), as well as findings *in vivo* (Figure 5), we tested whether spent media of virally infected host cells impacted bacterial growth or attachment in varying concentrations of bacitracin. *S. pneumoniae* growth in the presence of bacitracin was not significantly altered in spent media of uninfected or IAV-infected host cells (Figure 6D). This indicates that cells do not secrete a factor that is synthetically lethal with bacitracin. Intriguingly, while not significantly altering the IC_50_ (Figure S6C), we observed a combinatorial effect of spent media from infected cells with bacitracin on attachment to naïve host cells at specific concentrations (Figure 6E). This indicates that the infection-dependent effect of bacitracin treatment can in part be attributed to changes in the microenvironment of IAV-infected cells.

### Metabolic rewiring by IAV and *S. pneumoniae* shapes the infection niche and impacts pathogenicity

The cellular microenvironment and its reshaping during viral pre-infection is contributing to the pathogenicity of *S. pneumoniae*^63^. Indeed, we identified glycolytic enzymes, as well as regulators of purine- and pyrimidine synthesis as important for bacterial adherence and intracellular survival (Figure 2, Figure 3). To investigate the putative reshaping of the microenvironment during infection, we performed untargeted metabolomics on supernatants collected from A549 cells during viral, bacterial and co-infection (Figure S7A). Although most detected metabolites remained unchanged, 33 metabolites (29.2%) displayed substantial alterations relative to uninfected controls (Figure 7A). Viral infection alone caused only minor changes, whereas pneumococcal infection strongly remodeled the extracellular metabolite landscape. The most prominent differences involved nucleosides and their derivatives.

**Figure 7:**
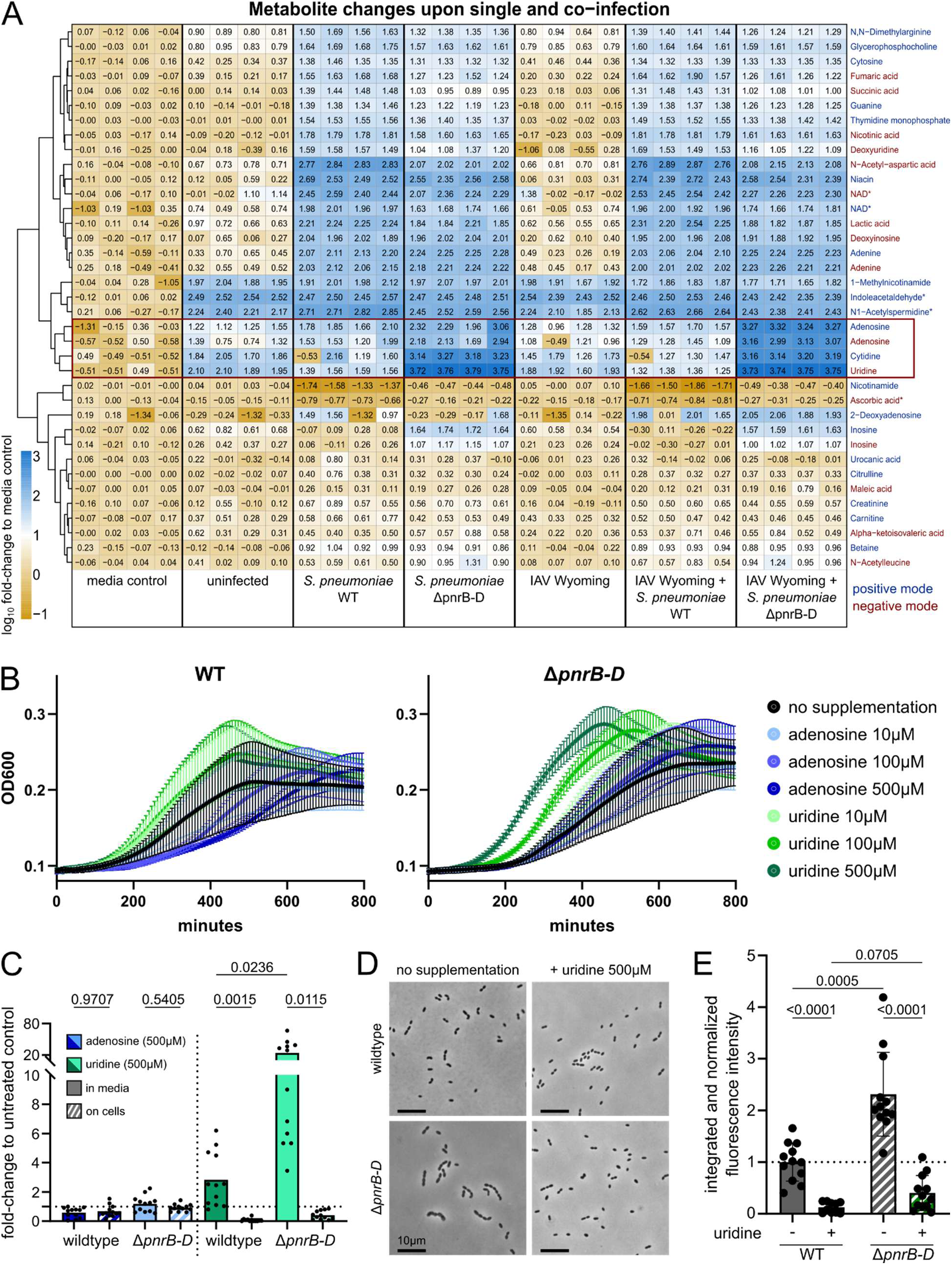
Metabolomics and the role of purine / pyrimidine biosynthesis. A) Heatmap of all metabolites that change during single and co-infection (at least 5-fold enrichment or depletion in at least one condition) with respect to the media control. Metabolites are indicated by name and whether they were identified in positive mode (labeled in blue) or in negative mode (labeled in red). The red box indicates nucleosides that were most strongly altered during infection, and that are further characterized. B) Growth curves (OD_600_) of wildtype (left) and *ΔpnrB-D* (right) *S. pneumoniae* in OptiMEM media without supplementation (black curves), supplemented with adenosine (shades of blue) or uridine (shades of green). Error bars indicate standard deviation across three biological replicates. C) CFUs recovered in media (solid bars) and after host cell attachment (striped bars) in the presence of adenosine (blue) or uridine (green). All values are normalized to the untreated control. P-values were calculated by one-way ANOVA. D) Light microscopy images of *S. pneumoniae* wildtype (top) or *ΔpnrB-D* mutant (bottom) with (right) or without (left) uridine supplementation. Representative images across three biological replicates and at least 3 technical replicates. Scale bar: 10μm. E) Quantification of fluorescence microscopy (example image displayed in Figure S7) after staining with Serotype-II antibody. Capsule expression was quantified and normalized to cell number for *S. pneumoniae* wildtype or *ΔpnrB-D* mutant in the presence or absence of uridine as indicated. Bars represent mean with standard deviation across three biological replicates, at least three images acquired per replicate, and p-values were calculated by one-way ANOVA.

To further investigate nucleotide metabolism, we compared wildtype *S. pneumoniae* with a Δ*pnrB-D* mutant lacking components of the Pnr nucleoside transporter^64,65^. This mutant displayed increased attachment to A549 cells (Figure 2), yet has a moderate growth deficit. Several nucleosides, including adenosine, uridine and cytidine, accumulated in supernatants infected with the mutant strain (Figure 7A, red box), consistent with impaired metabolite uptake.

Next, we tested whether altered nucleoside availability affected bacterial growth. Adenosine supplementation reduced growth of the wildtype strain but had minimal impact on Δ*pnrB-D*, whereas uridine promoted growth in both strains in a concentration-dependent manner (Figure 7B). Notably, uridine supplementation largely rescued the growth defect of Δ*pnrB-D* (Figure S7B-C). Other tested metabolites produced comparatively minor effects or increased the growth for both strains equally (Figure S7D).

We then examined whether nucleosides influenced host-cell attachment. Adenosine did not affect attachment beyond its effects on bacterial growth (Figure 7C, blue bars). In contrast, uridine increased bacterial growth while simultaneously reducing host-cell attachment (Figure 7C, green bars). This phenotype could not be attributed to host-cell toxicity (Figure S7E) or host-cell preconditioning by uridine (Figure S7F), suggesting a direct effect on bacterial physiology or morphology.

To identify underlying mechanisms, we assessed bacterial morphology and capsule production. The Δ*pnrB-D* mutant formed longer chains than the wildtype strain, a phenotype that was reversed by uridine supplementation (Figure 7D). *S. pneumoniae* growth behavior has previously been linked to capsule expression and composition^66,67^. Antibody staining for capsule expression revealed that the *ΔpnrB-D* mutant produces more capsule (Figure S7G) compared to the wildtype D39V strain. During uridine supplementation, this effect could be reversed, and capsule expression was significantly lower compared to the untreated control in both wildtype and mutant (Figure 7E). This is in line with published work linking nucleoside metabolism to capsule expression and pathogenicity^68^.

Together, these findings identify nucleotide metabolism, and particularly uridine availability, as an important regulator of pneumococcal physiology during infection. Alterations in nucleoside utilization influence growth, morphology, capsule production and host-cell attachment, revealing a trade-off between bacterial proliferation and adherence.

## Discussion

In this work, we systematically identify *S. pneumoniae* genes that contribute to epithelial attachment and intracellular survival during both single infection and IAV co-infection using genome-wide CRISPRi screening. Overall, only a small fraction of genes displayed condition-specific fitness effects, highlighting the robustness of core pneumococcal pathogenicity determinants across infection contexts. Nevertheless, several genes involved in cell envelope biogenesis and nucleotide metabolism exhibited altered importance during co-infection, indicating that influenza infection modifies specific bacterial vulnerabilities rather than globally rewiring pneumococcal fitness requirements.

We highlight the importance of cell wall-associated processes during infection. Multiple genes involved in peptidoglycan and cell envelope biogenesis contributed to host-cell attachment, and both genetic and pharmacological perturbation converged on these pathways. In particular, depletion of *uppP* strongly impaired attachment specifically during IAV co-infection. UppP-mediated undecaprenyl phosphate recycling is essential for efficient cell wall synthesis and has previously been linked to pneumococcal virulence *in vivo*^50^, supporting the biological relevance of our findings. Together with recent work highlighting cell wall homeostasis as a central determinant of antibiotic susceptibility and host adaptation in pneumococci^69–72^, our data suggest that envelope biosynthesis remains an attractive target for anti-infective intervention, particularly in the context of viral-bacterial co-infections.

Comparison with published CRISPRi datasets and *in vivo* studies^49^ revealed only modest overlap at the level of individual genes. While this initially appears surprising, it likely reflects the highly context-dependent nature of pneumococcal gene fitness. Previous studies have demonstrated substantial variation in genetic requirements across strains, host niches, and environmental conditions, indicating that pathogenicity cannot be explained by a universal set of virulence genes alone^30,31^. Our results therefore reinforce the concept that bacterial fitness emerges from the interaction between genotype and infection environment. In this regard, reductionist *in vitro* systems complement *in vivo* models by enabling mechanistic dissection of host-pathogen interactions with substantially greater throughput.

Integration of metabolomics with functional genetics highlighted nucleotide metabolism as a second major determinant of pathogenicity during co-infection, which is in line with recent studies^73–76^. Pneumococcal infection caused marked changes in the level of extracellular purines and pyrimidines, whereas viral infection alone had a comparatively minor effect on metabolite abundance. These findings are consistent with previous studies showing that *S. pneumoniae* relies heavily on nucleoside acquisition and salvage pathways during growth and infection^77^. Several genes involved in nucleotide transport and regulation were identified in our screen, and deletion of the nucleoside transporter operon *pnrB-D* altered both extracellular metabolite profiles and infection phenotypes. This complements a recent study on *pnrA*, linking the protein to iron homeostasis^65^. Recent work has similarly demonstrated that uracil transport and pyrimidine metabolism, including NAD^+^ production contribute to pneumococcal fitness and virulence^68,78,79^. Additionally, carbon source variation has been shown to fine-tune capsule production, thickness and pathogenicity^80^. Finally, given the centrality of nucleotide and nucleoside metabolism, modulations in and perturbations of the identified pathways also has implications beyond infection, such as affecting biofilm formation^81^ or antimicrobial resistance^72,82^.

Our data suggests a mechanistic link between uridine availability, capsule regulation, and bacterial morphology. Pyrimidine metabolism has previously been connected to capsule biosynthesis, and uridine triphosphate (UTP) is a required precursor for polysaccharide capsule production^83^. Capsule abundance alone does not readily explain the observed decrease in host-cell attachment, since unencapsulated pneumococci generally adhere more efficiently to epithelial cells^84^. Rather than being linked to changed capsule production or composition, increased attachment in uridine depletion (e.g. in the *ΔpnrB-D* mutant) likely originates from changes in chain length, which has been shown to promote pneumococcal adherence and colonization^85^. This is in line with several chain-length altering genes increasing attachment in the screening^86^. Additionally, chain length has also been implicated in competence^87^, which is another determinant of pathogenicity, and should therefore be investigated further.

Despite the novel insights, several limitations of this study remain: First, the cell culture infection models, while allowing for higher throughput, cannot fully recapitulate the complexity of the respiratory tract, where epithelial heterogeneity, tissue architecture, immune cell recruitment, and microbiota-derived factors contribute to disease progression. Several new culturing methods, including air-liquid interface systems and co-cultures^88–90^ can help to link our findings to a more physiological context. Depletion of protein levels by CRISPRi also requires several bacterial generations and might not be limiting enough under our experimental conditions. Second, differences between influenza strains suggest that virus-specific effects on host physiology may influence bacterial fitness landscapes. Finally, the screening was performed in a defined pneumococcal background, whereas pneumococcal virulence traits, particularly capsule regulation, vary substantially across strains and serotypes^91^. Future studies should therefore investigate these mechanisms in primary cell systems and animal models while extending analyses to genetically diverse pneumococcal isolates.

In conclusion, our study provides a genome-scale resource of pneumococcal fitness determinants during host-cell attachment and intracellular survival. The data identify cell envelope homeostasis and nucleoside metabolism as major regulators of pathogenicity and reveal context-specific vulnerabilities that emerge during influenza co-infection. These findings highlight a finely tuned balance between growth, capsule production and pathogenicity. The identification of these molecular interaction points between the host and co-occurring pathogens improves our understanding of pneumococcal adaptation to the host environment and highlights potential targets for future antimicrobial and anti-infective strategies.

## Supplementary material

**Figure S1:**
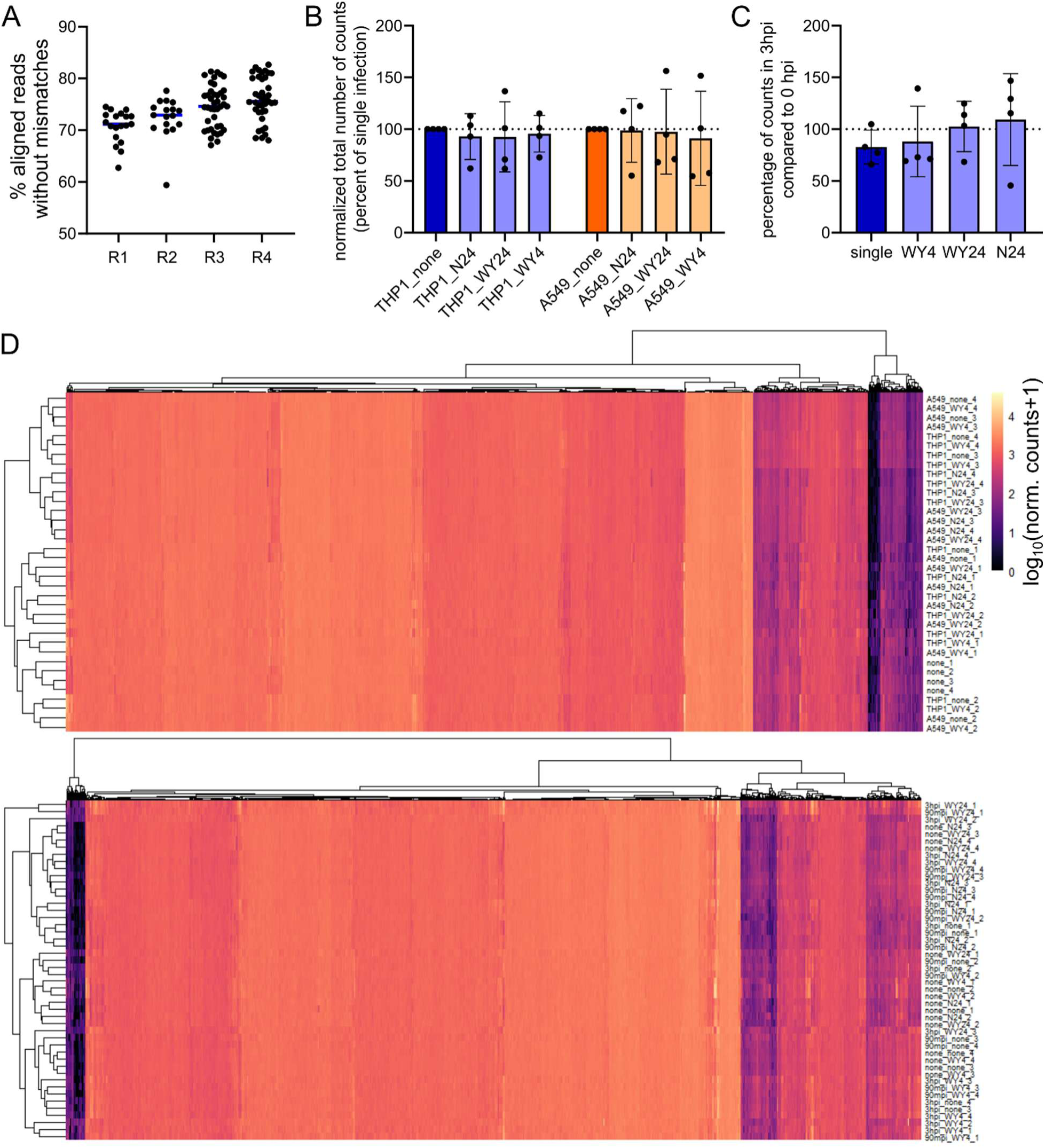
In depth quality control of CRISPRi screening data. A) Percentage of aligned reads in each of the four replicates. Each dot represents an infection condition (cell line, viral pre-infection) and time point. B) Total number of sgRNA counts (sum of all) normalized to the single infection control. This bar graph shows that co-infection does not significantly alter overall read counts. C) As in panel B, but for intracellular survival. Total number of reads is not changed at later time points. D) Heatmaps depicting the normalized number of reads (including pseudo-counts) for attachment (top) and intracellular survival (bottom). Only a minority of genes displayed low counts, showing that bottlenecks are not a major problem for the study.

**Figure S2:**
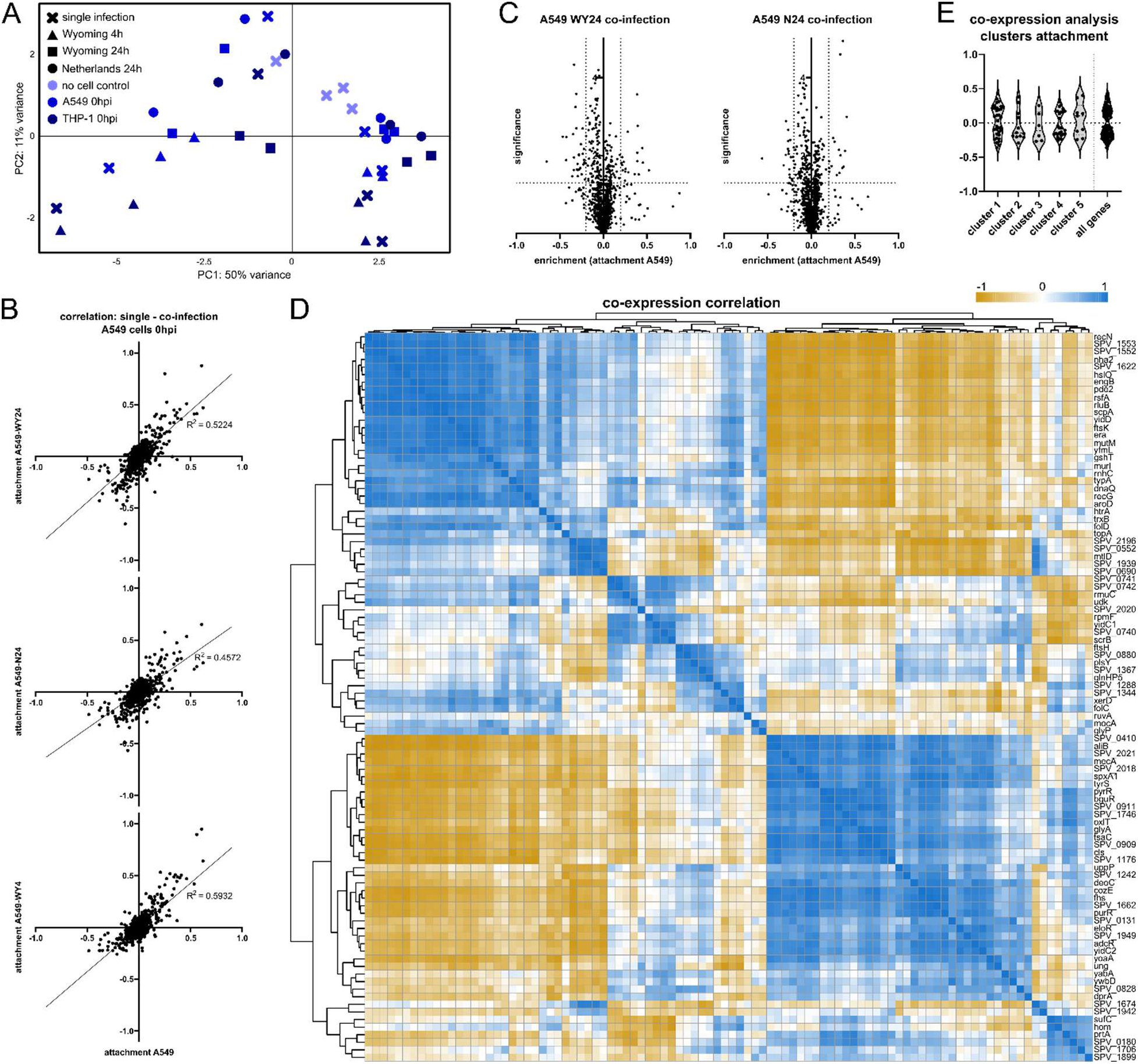
in-depth assessment of essentiality during attachment. A) PCA plot displaying global variance in different viral co-infection conditions and cell lines. B) Correlation in attachment scores between single infection (x-axis) and different co-infection conditions (y-axes). Line depicts linear regression and Pearson correlation coefficient is indicated. C) Volcano plots (as in Figure 2C), but for the remaining co-infection conditions (24h pre-infection). D) Co-expression correlation of all genes that altered attachment when targeted by CRISPRi: Co-expression with all *S. pneumoniae* genes^42^ was determined and transformed into a co-expression fingerprint. Pairwise correlations were calculated: Perfect correlation (score = 1) is colored in blue, anti-correlation (score = −1) in orange. D) K-means clustering (k=5) was performed to group candidate genes into clusters with similar fingerprints and distribution of z-scores in the screening is displayed as violin plots for each cluster.

**Figure S3:**
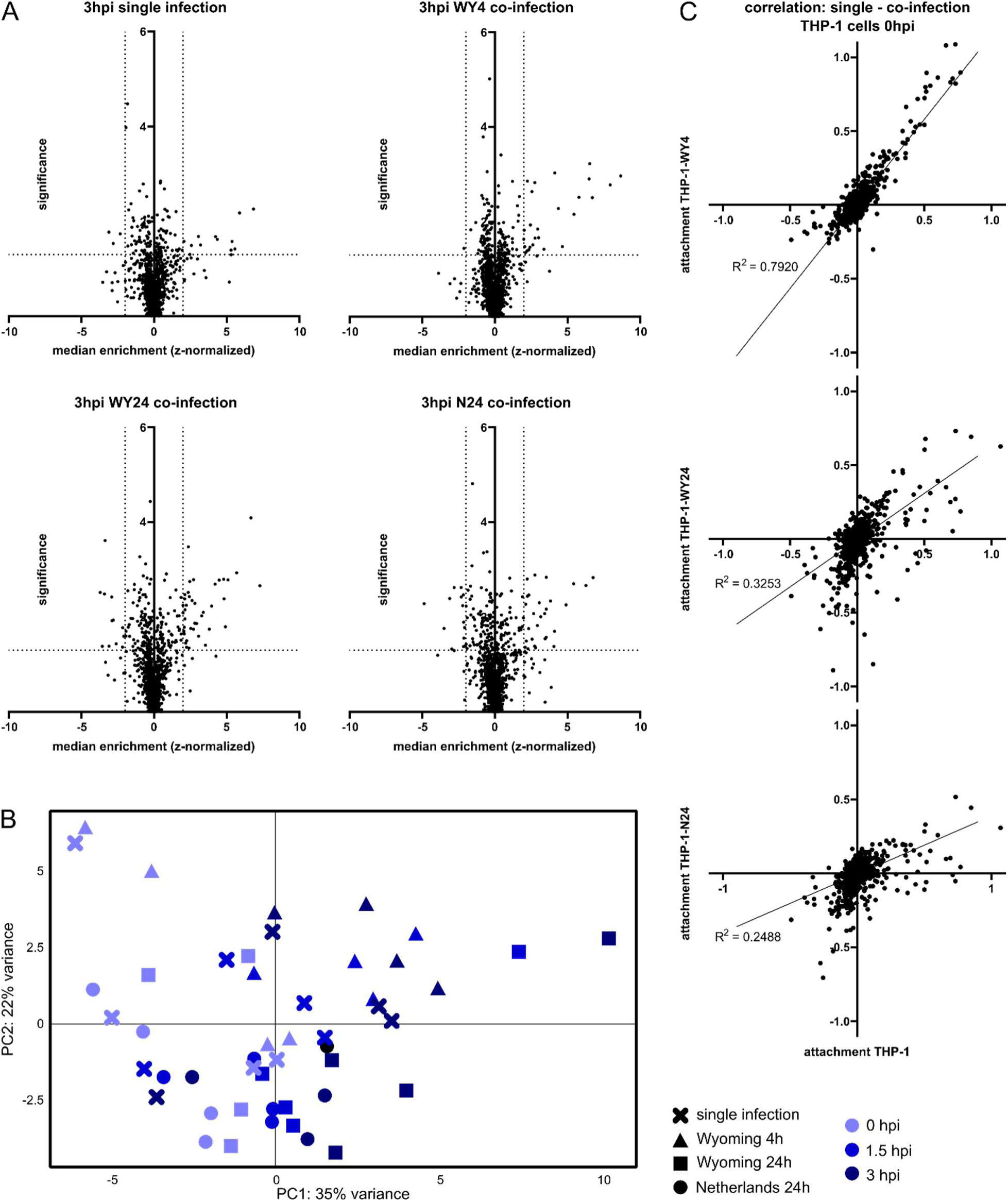
In-depth assessment of genes altering intracellular survival in CRISPRi screening. A) Volcano plots depicting the enrichments for intracellular survival during single and co-infection. X-axis indicates the z-score for each gene, the y-axis the negative logarithmic FDR. Each dot represents one gene. B) PCA of enrichment fingerprints for each of the assessed conditions and timepoints as indicated. While variance between the samples is comparably small, there is an observable shift from early (lighter blue) to later (darker blue) timepoints. C) Correlations of enrichment scores (log-fold changes) in co-infection (y-axes) compared to single infection (x-axis) for THP-1 cells at timepoint 0. Pearson correlation, as well as linear regression line are indicated.

**Figure S4:**
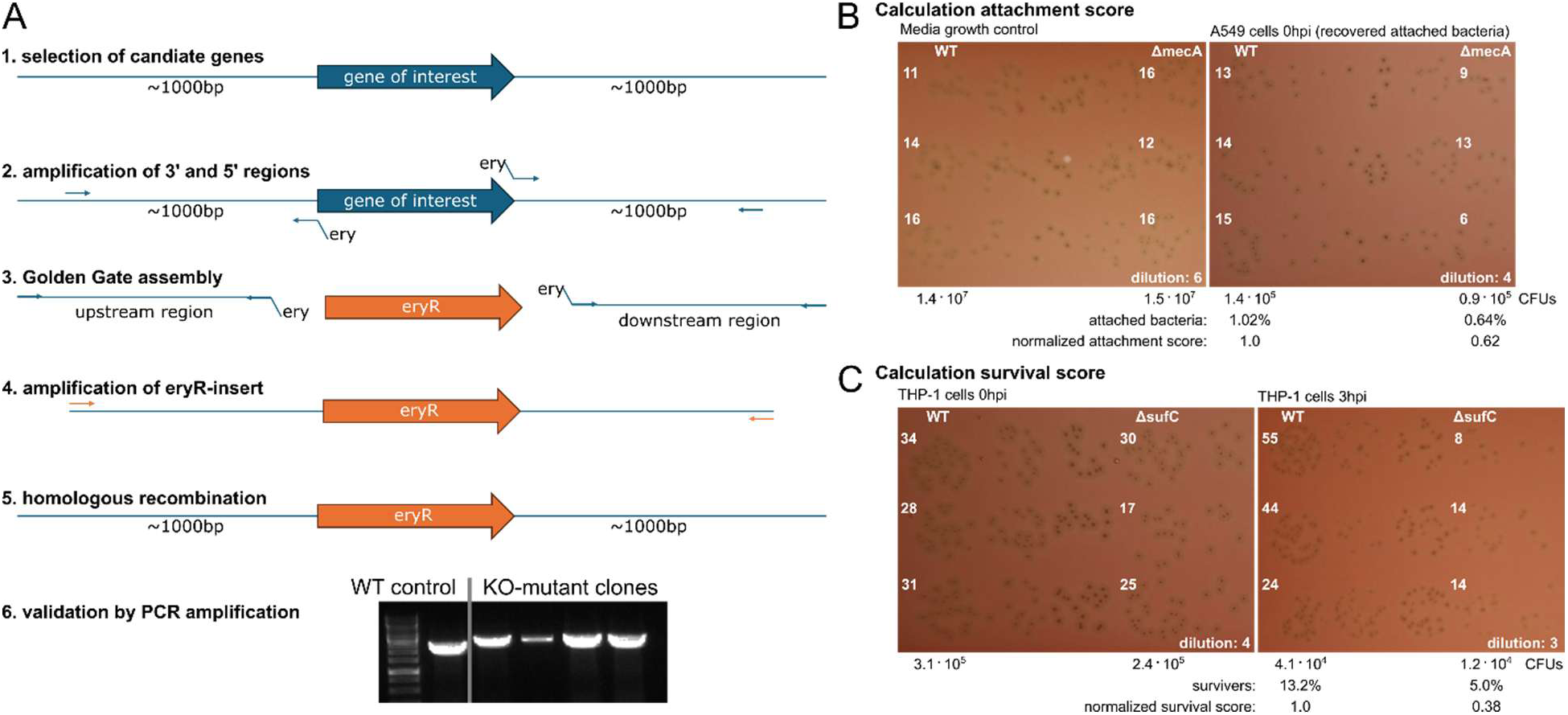
Validation strategy and calculation. A) Workflow for the creation of clean deletion mutants as described in the Methodology section. In brief, candidate genes were selected and upstream and downstream regions were amplified using overhangs to an erythromycin cassette. New inserts were created by Golden Gate Assembly, amplified by PCR and subsequently inserted into the genome by recombination. Finally, the mutant was confirmed by PCR. B) Calculation example for attachment scores. CFUs were quantified for media growth, as well as bacteria recovered after attachment for each mutant, as well as the wildtype. The fraction of attached bacteria was calculated (with respect to the CFUs recovered from media), and subsequently these were normalized to WT. C) Similarly to panel B, CFUs were quantified at 0hpi and 3hpi for all mutants and the wildtype. The fraction of surviving bacteria was calculated and then normalized to the percentage in WT infection.

**Figure S5.**
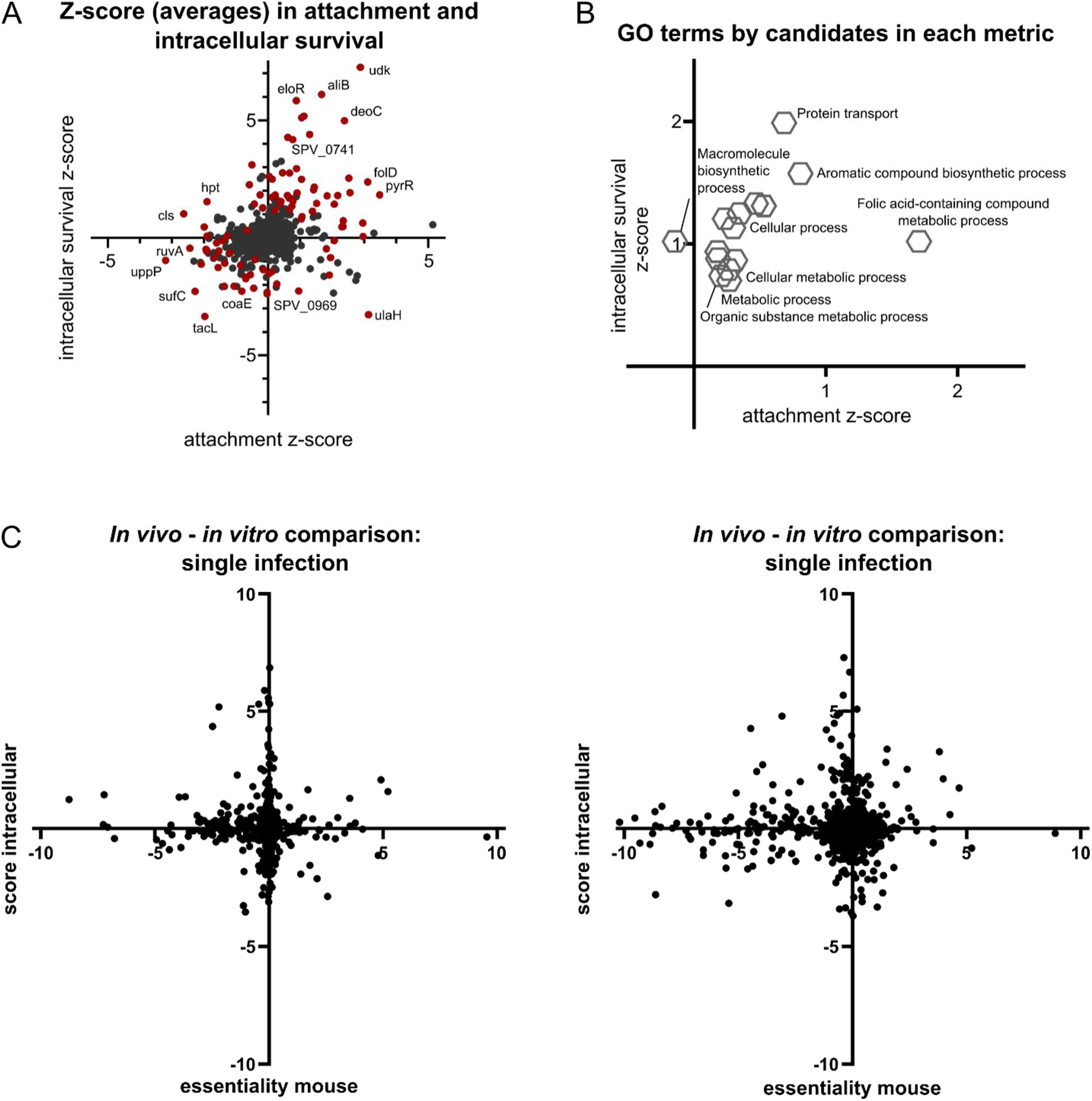
Global analysis, GO-terms and additional comparison graphs. A) Correlation of intracellular survival score (averages across infection conditions, y-axis) to attachment scores (averages across infection conditions, x-axis). Hits are indicated in red, and several are indicated by name (subset chosen for visibility). B) Scatterplot of GO-terms (shown in Figure 5A) by enrichments (z-scores) of the genes each term is comprised of. For each term, the genes that were associated with it were taken as basis to calculate the mean z-score in attachment (x-axis) and intracellular survival (y-axis). C) Scatterplots of the comparison of *in vivo* CRISPRi study^49^ (x-axis) to intracellular survival score in single (left) or co-infection (right). No significant correlation could be detected.

**Figure S6:**
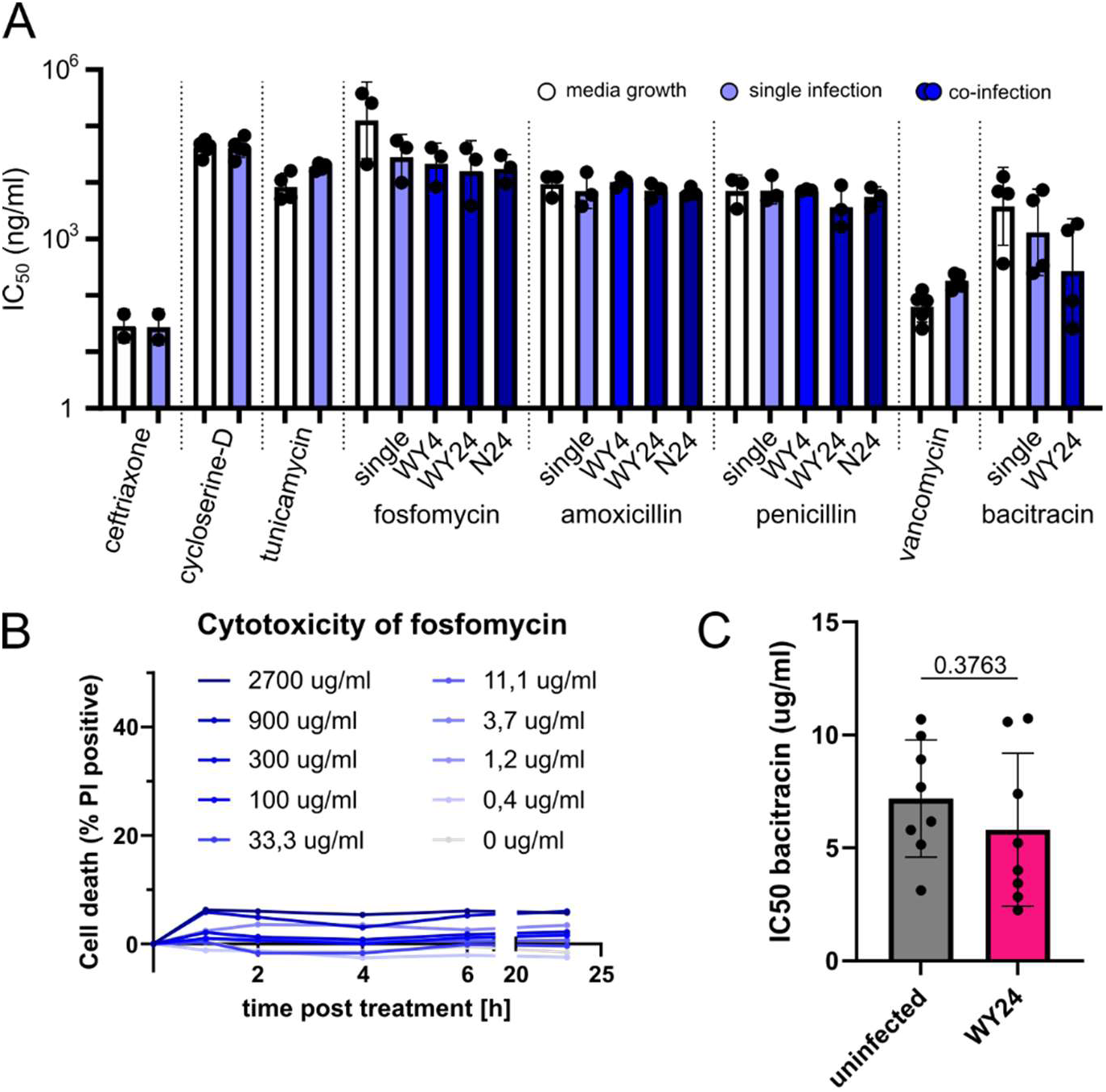
Impact of antibiotics on *S. pneumoniae* and host cells. A) All calculated IC_50_ values in media (white bars) and for attachment (blue bars) for each of the tested antibiotics. CFU counts across a range of antibiotic concentrations was used as basis for the calculation. Each dot represents a biological replicate, each of which is comprised of at least three technical replicates. B) Host cell cytotoxicity (by propidium iodide staining) over time in different concentrations of fosfomycin. Cell death was normalized to 0% at the beginning of the treatment. C) Calculation of the IC_50_ for attachment in spent media of uninfected (grey) or IAV-infected cells (pink). CFUs were quantified across a range of bacitracin concentrations to calculate IC_50_. Each dot represents a replicate across three independent infection experiments.

**Figure S7:**
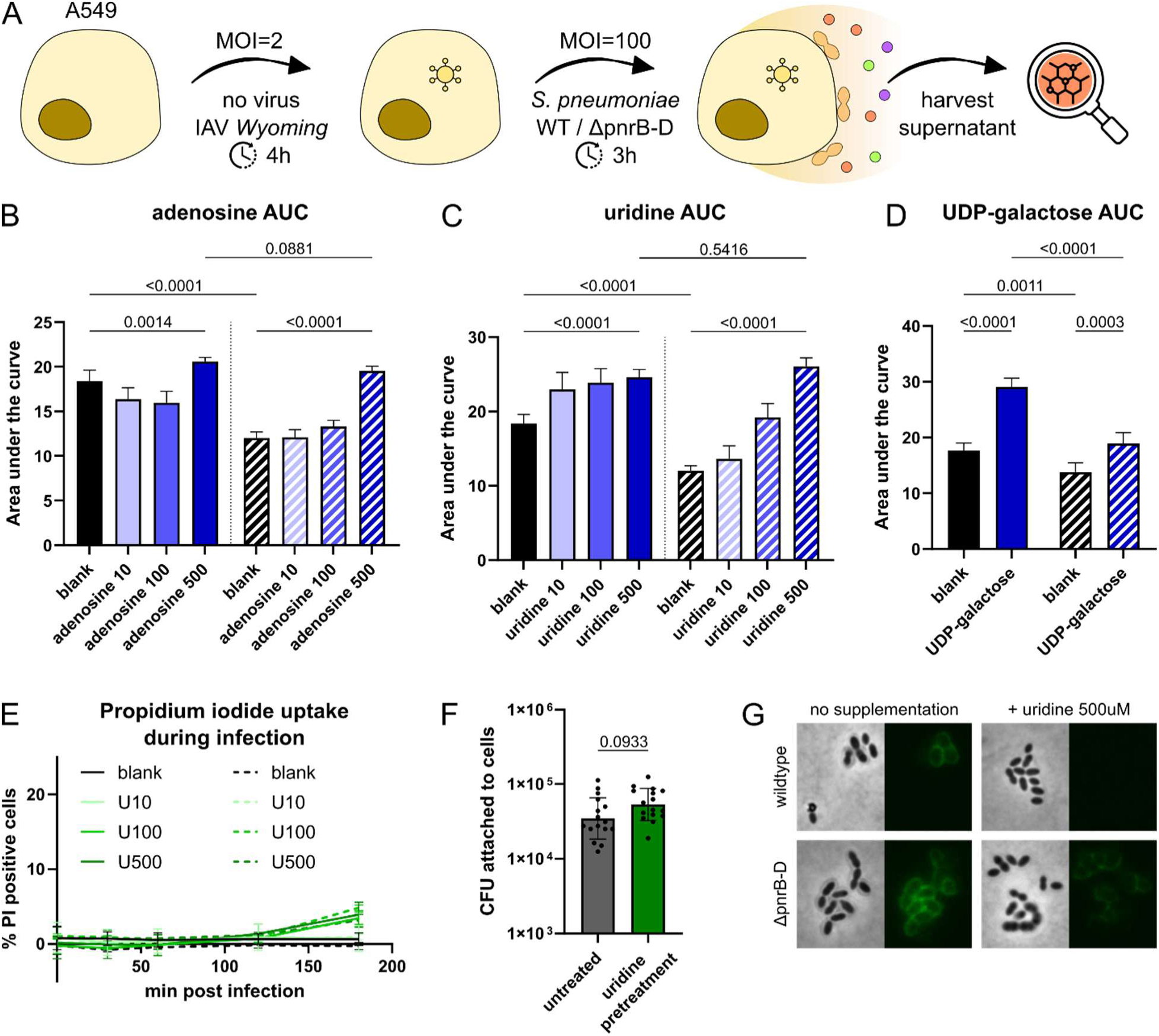
Nucleosides change growth and pathogenicity of *S. pneumoniae*. A) Workflow for untargeted metabolomics during *in vitro* infection with IAV and / or *S. pneumoniae*. A549 cells were infected for 4h with the *Wyoming* strain (uninfected control included), and subsequently challenged with *S. pneumoniae* (wildtype, *ΔpnrB-D* mutant or uninfected control). Cells were spun down, supernatant was harvested and analyzed in positive and negative mode and compared to metabolite profiles of a blank media control (which was never exposed to cells). B-D) Bar charts quantifying the area under the curve of OD_600_ growth curves (up to 12h of measurements) of wildtype or *ΔpnrB-D* mutant *S. pneumoniae* during supplementation with the indicated metabolite at different concentrations: adenosine (panel B), uridine (panel C) and UDP-galactose (panel D). Bars depict mean and standard deviation across at least three biological replicates. E) Cytotoxicity assay (PI uptake) of A549 cell exposed to uridine at different concentrations during infection with wildtype (solid lines) or *ΔpnrB-D* mutant *S. pneumoniae* (dashed lines). F) Quantification of CFUs attached to A549 cells that were treated for 5h with uridine (500μM) prior to infection with *S. pneumoniae* (wildtype) in the absence of uridine. Each dot represents a replicate across three independent experiments. G) Brightfield and fluorescence microscopy of wildtype (top) and *ΔpnrB-D* mutant (bottom) *S. pneumoniae* after treatment with uridine (500μM) for 2.5h and subsequent antibody staining for capsule. This is a representative image, and quantification of several fields of view across biological replicates is depicted in Figure 7E.

## Materials and methods

### Ethics statement

The research described in this study complies with all relevant ethical regulations. No animals or animal-derived cells were used as part of this work.

### Data availability

Source data has been made available in Mendeley Data (DOI 10.17632/xjdkgj22tn.1). This includes original images and full datasets.. All research material, analysis algorithms or additional data are available upon request.

### Statistical analysis

Analyses, significance testing and visualization, unless indicated otherwise, was performed in Prism (version 11.0.1). Necessary prerequisites for test selection (multiple testing, Welsh correction, sample pairedness, normalization) were taken into account. Further details of statistical analysis are indicated where required.

### Bacterial strains, viral strains and cell culture

*Streptococcus pneumoniae* strain D39V was used throughout this study. For genome-wide fitness screening, an IPTG-inducible CRISPR interference (CRISPRi) library constructed in a capsule-deficient (*Δcps*) D39V background was employed. To do so, the previously published library^49^ was transformed into a markerless *Δcps* background^92^. Pneumococci were routinely cultured in C+Y medium^93^ under standard laboratory conditions. For validation experiments, clean deletion mutants were generated in the encapsulated D39V wild-type background as described below.

Influenza A virus strains A/Netherlands/1005/2009 (H1N1; hereafter referred to as Netherlands) and Wyoming (H3N2) were used for co-infection experiments. Viral stocks were kindly provided by Prof. Mirko Schmolke (University of Geneva, Switzerland). Cells were infected with IAV at a multiplicity of infection (MOI) of 2 either 4h or 24h prior to bacterial challenge, as specified for individual experiments.

Human alveolar epithelial A549 cells were maintained in Dulbecco’s Modified Eagle Medium (DMEM; GlutaMAX formulation, gibco, ref. 31966-021) supplemented with 10% fetal bovine serum (FBS, Thermo Fisher, ref. 17593595). Cells were cultured at 37°C in a humidified atmosphere containing 5% CO₂ and were used up to passage 15. THP-1 human monocytic cells were maintained in Roswell Park Memorial Institute (RPMI) 1640 medium with GlutaMAX (gibco, 61870-010) supplemented with 10% FBS under standard culture conditions (37°C, 5% CO₂).

For generation of macrophage-like cells, THP-1 cells were seeded into the appropriate culture vessels 48 h prior to infection and differentiated using phorbol 12-myristate 13-acetate (PMA, Sigma Aldrich, P8139-1MG). Following differentiation, cells were used for bacterial infection experiments and intracellular survival assays.

### Library preparation for CRISPRi during infection

The pooled CRISPRi library was routinely propagated in C+Y medium at 37°C. Prior to infection experiments, the library was pre-induced with 40μM IPTG and cultured for approximately 12 bacterial generations to allow depletion of target gene products and establishment of the knockdown phenotype. Following pre-induction, library aliquots were stored as glycerol stocks and used as the inoculum source for all screening experiments.

For each infection experiment, an aliquot of the pre-induced glycerol stock was inoculated into fresh C+Y medium supplemented with IPTG and incubated at 37°C. Cultures were grown for 3 to 3.5h to mid-exponential phase (OD_600_ ≈ 0.2) before being used for infection assays.

### Viral infection

For IAV infection, cell monolayers were washed once with phosphate-buffered saline (PBS, prepared in-house) prior to inoculation. Cells were infected with 100µl of virus suspension in OptiMEM (gibco, ref. 31985-070) at MOI = 2 and incubated for 1h at 37°C in a humidified atmosphere containing 5% CO₂ to allow viral adsorption.

Following incubation, the viral inoculum was removed, and cells were washed once with PBS to eliminate unbound virus particles. Subsequently, 100µL OptiMEM supplemented with TPCK-treated trypsin (Merck, T1426, 1:2000 dilution of a freshly thawed 2mg/ml stock for a final concentration of 1µg/ml) was added to each well. Infected cells were then incubated for either 4h or 24h at 37°C and 5% CO₂ prior to secondary bacterial infection, as specified for the respective experiment. Mock-infected controls were treated identically but received virus-free medium during the adsorption step.

### Bacterial infection

*S. pneumoniae* cultures were prepared as described above and grown to an OD_600_ of 0.2. Bacteria were collected by centrifugation, washed once with phosphate-buffered saline (PBS), and resuspended in OptiMEM. Host cells were infected at MOI = 100 and incubated at 37°C in a humidified atmosphere containing 5% CO₂. After 3h of incubation, corresponding to time point 0 (T0), samples were processed for downstream analyses. As a media control, bacteria were incubated under identical conditions in OptiMEM in the absence of host cells.

For determination of bacterial attachment, infected monolayers were washed three times with PBS to remove non-adherent bacteria. Cells were subsequently lysed with 0.1% Triton X-100 (ITW Reagents), and the recovered lysates were collected for quantification of cell-associated bacteria. For intracellular survival assays, infected monolayers were washed three times with PBS after the initial 3h infection period and the medium was replaced with fresh OptiMEM containing recombinant LytA (1μM) to eliminate extracellular bacteria. At 1.5h and 3h post-internalization, respectively, cells were washed three times with PBS and lysed with 0.1% Triton X-100.

### CRISPRi-seq: DNA extraction, library preparation, sequencing and analysis

Genomic DNA was isolated from bacterial pellets that were recovered after infection experiments using the Vazyme FastPure Bacteria DNA Isolation Mini Kit (DC103-01), following the instructions by the manufacturer. After extraction DNA was resuspended in molecular-grade water, and stored at −20 °C.

For CRISPRi library analysis, sgRNA cassettes were amplified from genomic DNA by PCR using indexed primers annealing to sequences flanking the sgRNA region. Sample-specific barcodes were incorporated during amplification to enable multiplexed sequencing. Library preparation was performed according to the CRISPRi-seq workflow described previously by de Bakker *et al.*^25^.

Indexed amplicons were pooled and prepared for sequencing according to the manufacturer’s recommendations and sequenced on an AVITI sequencing platform using short-read sequencing. sgRNA abundances were quantified from sequencing reads using 2FAST2Q^94^, and differential abundance analyses were performed in R-studio (version 2025.09.1 build 401) running with R (v4.5.1) using DESeq2^95^. A negative binomial generalized linear model was used to estimate log₂ fold changes and significance, with *P* values adjusted for multiple testing using the Benjamini-Hochberg false discovery rate (FDR) method. The analysis pipeline is available on Mendeley Data (DOI: 10.17632/xjdkgj22tn.1).

For visualization, log₂ fold change values were scaled on a gene-wise basis without centering prior to heatmap generation, enabling comparison of relative fitness effects across conditions while preserving directionality of enrichment and depletion. Principal component analysis, correlation analysis, and data visualization were performed in R and GraphPad prism (version 11.0.1).

### Creation of clean deletion mutants

Clean deletion mutants were generated by Golden Gate assembly and allelic replacement in *S. pneumoniae* D39V as outlined in Figure S4A. Briefly, approximately 1kb genomic regions upstream and downstream of the target gene were amplified by PCR using primers containing overhangs complementary to an erythromycin resistance cassette. The upstream fragment, erythromycin resistance cassette, and downstream fragment were assembled by Golden Gate cloning to generate a deletion construct.

Assembly reactions were analyzed by agarose gel electrophoresis, and correctly assembled products were subsequently amplified by PCR to obtain sufficient quantities of linear DNA for transformation and to enhance homologous recombination efficiency. The resulting PCR products were introduced into competent pneumococci by natural transformation. Transformants were selected on Columbia blood agar plates (prepared in-house) supplemented with erythromycin and re-streaked three consecutive times on selective medium to ensure clonality. Successful deletion of the target locus was confirmed by colony PCR using primers flanking the recombination region. Expected amplicon sizes were verified by agarose gel electrophoresis before mutants were used for subsequent experiments.

### Treatment with antibiotics or nucleosides during infection

Infection experiments in the presence of antibiotics or nucleosides were performed as described above for the respective infection assays. To ensure that the desired final compound concentration and MOI were reached upon addition to host cells, antibiotic- or nucleoside-containing media were prepared at 2-times the intended final concentration. Likewise, bacterial suspensions were prepared at 2-times the desired MOI. Equal volumes of compound-containing medium and bacterial suspension were mixed immediately prior to infection, and half of the resulting volume was added to host cells. The remaining volume was incubated under identical conditions in the absence of host cells and served as a media control.

For determination of inhibitory concentrations (IC_50_), bacterial burden was quantified by CFU enumeration following infection. Dose-response curves were generated using GraphPad Prism version 11.0.1 by fitting the data with the “inhibitor versus response” nonlinear regression model using three parameters. For visualization of combined datasets, CFU values were normalized using the upper and lower CFU boundaries within each biological replicate. IC_50_ values were calculated independently for each biological replicate, and the resulting values were used for subsequent statistical analyses.

To determine cytotoxicity, propidium iodide (Merck, P4864-10ml) uptake assay was conducted. Cell permeabilization was quantified with respect to total lysis control (Tx-100 treated) after determining fluorescence on a Tecan spark plate reader using the appropriate fluorescence filters in the spark control software (version 3.2).

### Untargeted metabolomics during infection

LC-MS grade water, acetonitrile (ACN) and methanol (MeOH) were obtained from Th. Geyer (Germany). High-purity ammonium acetate, ammonium formate, ammonium hydroxide, and formic acid were purchased from Merck (Germany). Stable isotope labelled amino acids (MSK-A2-1.2; Cambridge Isotope Laboratories, MA, USA) were used as internal standards for untargeted metabolomics at a concentration of 0.5 % in the final sample.

Infections were performed as described above. Conditioned cell culture supernatants were collected from the following experimental conditions: (i) OptiMEM medium control not exposed to cells, bacteria, or virus; (ii) uninfected cells; (iii) cells infected with IAV; (iv) cells infected with *S. pneumoniae* wild type; (v) cells infected with the *S. pneumoniae* Δ*pnrB-D* mutant strain; (vi) cells co-infected with IAV and *S. pneumoniae* wild type; and (vii) cells co-infected with IAV and the Δ*pnrB-D* mutant strain. All infections were performed in OptiMEM, and cell culture supernatants were collected at the indicated time points and stored at −80°C until analysis.

80µL of cell culture supernatant were extracted via addition of 320µL of MeOH:ACN (1:1, v:v; including internal standards). After shaking 4°C at 1800rpm for 30 min and incubation at −20°C for 30 min, the samples were centrifuged at 15,000 × g and 4°C for 10 min using a 5415R microcentrifuge (Eppendorf, Hamburg, Germany). Extracts were dried in a purged nitrogen environment using a Genevac EZ-Elite 2.4 vacuum evaporator (ATS Life Sciences, GA, USA). Dried samples were reconstituted in 80µL 80% MeOH, centrifuged and transferred to analytical glass vials for LC-MS/MS analysis. The analysis was initiated within one hour after completion of sample preparation.

LC-MS/MS analysis was performed on a Vanquish UHPLC system coupled to an Orbitrap Exploris 240 high-resolution mass spectrometer (Thermo Fisher Scientific, MA, USA) in negative and positive ESI (electrospray ionization) mode. Chromatographic separation was carried out on an Atlantis Premier BEH Z-HILIC column (Waters, MA, USA; 2.1 mm x 100 mm, 1.7µm) at a flow rate of 0.25 mL/min. The mobile phase consisted of water:acetonitrile (9:1, v/v; mobile phase phase A) and acetonitrile:water (9:1, v/v; mobile phase B), which were modified with a total buffer concentration of 10mM ammonium acetate (negative mode) and 10mM ammonium formate (positive mode), respectively. The aqueous portion of each mobile phase was pH-adjusted (negative mode: pH 9.0 via addition of ammonium hydroxide; positive mode: pH 3.0 via addition of formic acid). The following gradient (20 min total run time including re-equilibration) was applied (time [min]/%B): 0/95, 2/95, 14.5/60, 16/60, 16.5/95, 20/95. Column temperature was maintained at 40°C, the autosampler was set to 4°C and sample injection volume was 5µL. Analytes were recorded via a full scan with a mass resolving power of 120,000 over a mass range from 60 – 900 m/z (scan time: 100 ms, RF lens: 70%). To obtain MS/MS fragment spectra, data dependent acquisition was carried out (resolving power: 15,000; scan time: 22ms; stepped collision energies [%]: 30/50/70; cycle time: 900ms). Ion source parameters were set to the following values: spray voltage: 4100V (positive mode) / −3500V (negative mode), sheath gas: 30psi, auxiliary gas: 5psi, sweep gas: 0psi, ion transfer tube temperature: 350°C, vaporizer temperature: 300°C.

Untargeted metabolomics measurements were performed by the EMBL Metabolomics Core Facility (Heidelberg, Germany). All experimental samples were measured in a randomized manner. Pooled quality control (QC) samples were prepared by mixing equal aliquots from each processed sample. Multiple QCs were injected at the beginning of the analysis to equilibrate the analytical system. A QC sample was analyzed after every 5^th^ experimental sample to monitor instrument performance throughout the sequence. For determination of background signals and subsequent background filtering, an additional processed blank sample was recorded. Data was processed using MS-DIAL 4.9^96^ and raw peak areas were normalized via best-matched internal standard normalization (B-MIS)^97^ for relative metabolite quantification and downstream analyses. Level 1 feature identification was based on an in-house library for metabolomics (EMBL-MCF 2.0^98^) using accurate mass, isotope pattern, MS/MS fragmentation, and retention time information with a minimum matching score of 80%.

### Microscopy after uridine supplementation

For microscopy experiments, *S. pneumoniae* cultures were inoculated from glycerol stocks and grown in C+Y medium to an OD_600_ of 0.2. Cultures were subsequently diluted 1:10 into OptiMEM and incubated statically for 2.5h at 37°C in a humidified atmosphere containing 5% CO₂.

For immunofluorescence staining, bacterial cultures were incubated with primary antibody (Type-2 rabbit antiserum, SSI diagnostica, ref. 16745) at a 1:1000 dilution for 5min at 4°C. Cells were collected by centrifugation, washed once with ice-cold PBS, and resuspended in ice-cold PBS containing Alexa-488 coupled goat anti-rabbit secondary antibody (Invitrogen, A11008) at a 1:1000 dilution. Following incubation, bacteria were pelleted by centrifugation and resuspended in PBS at half of the original volume. Subsequently, 3µL of the stained bacterial suspension was spotted onto 1.2% agarose pads and imaged immediately.

Microscopy was performed using a Leica DMi 8 wide-field microscope equipped for brightfield and fluorescence imaging with a Leica DFC7000T camera. Brightfield images were acquired using an exposure time of 50ms, while fluorescence images were acquired using a GFP filter and 50ms exposure time. Images were acquired using a 100x objective with oil immersion (Type F immersion liquid, Leica). Image acquisition and initial processing were performed using Leica LAS X premium software (v3.1031329575). Further image analysis was carried out using ImageJ (version 1.53t).

For quantitative image analysis, bacterial masks were generated from the brightfield channel and used to determine the area of individual bacteria. For fluorescence quantification, background fluorescence was determined independently for each biological replicate and used to define a fluorescence threshold. Pixels exceeding the threshold were included in the analysis, and the integrated fluorescence intensity was calculated for each bacterium. To account for differences in cell size, integrated fluorescence values were normalized to the corresponding bacterial area.

## Acknowledgements

We would like to thank Mirko Schmolke, Mehdi Chabert-Ben Cherifa and Filomena Silva for providing the Influenza-A strains used in this study, alongside infection protocols and details on pathogen handling. We acknowledge the support of the EMBL Metabolomics Core Facility (MCF) in the acquisition and analysis of liquid chromatography-mass spectrometry data. Specifically, we would like to thank Dr. Bernhard Drotleff and Dr. Michael Zimmermann for their support.

## Author contributions

The study was designed by P.W. and J.W.V. All experiments and analysis were conceptualized and performed by P.W. Bacterial strains, unless cited, have been generated by P.W. under advice by J.W.V. Initial manuscript writing was done by P.W. and further refined by P.W., J.W.V. and P.B.

## Competing interests

The authors declare no competing interests.

## Materials & Correspondence

Request for materials and correspondence should be addressed to P.W. or J.W.V.

